# MRE11 suppresses germline mutagenesis at meiotic double-strand breaks in mice

**DOI:** 10.64898/2026.02.11.705388

**Authors:** Agnieszka Lukaszewicz, Thomas E. Wilson, Soonjoung Kim, Scott Keeney, Maria Jasin

## Abstract

SPO11 forms hundreds of double-strand breaks (DSBs) to initiate meiotic recombination that is normally error-free. However, SPO11 activity can be mutagenic when one chromatid incurs closely spaced DSBs (double cuts), especially when DSBs are dysregulated by loss of the ATM kinase. *De novo* indels and structural variants can arise via end joining at double cuts within a single hotspot (microdeletions) or at adjacent hotspots separated by at least 30 kb, as we now show, sometimes accompanied by ectopic insertions of double-cut fragments. Here, we investigate how meiotic DSB end processing influences end joining. In MRE11-deficient mouse spermatocytes, which do not resect their DSBs, deletions at double cuts occur readily, with end-joining breakpoint profiles closely matching SPO11 DSB profiles. Microdeletions suggest that two DSBs can be as close as ∼21 bp. The tyrosyl DNA phosphodiesterase TDP2 contributes to both deletion formation and ectopic insertion of double-cut fragments, presumably by removing SPO11 from DNA ends prior to joining. Finally, observations suggest a cooperative role for MRE11 and ATM in locally regulating DSB distributions. Our findings provide insight into the mechanism of *de novo* mutation origin, emphasizing the role of meiotic DSBs in shaping genome evolution.

## Introduction

Repair of meiotic double-strand breaks (DSBs) by homologous recombination ensures correct chromosome segregation and increases genetic diversity^1^. By contrast, end-joining pathways known to operate in mitotic cells^2^ are non-productive for the meiotic program^3^. Instead, end joining between closely-spaced meiotic DSBs (double cuts) can lead to genomic deletions and other mutagenic events in the mouse germline^4^. These events are detected in wild type but are much more frequent in the absence of the ATM kinase^4^. How cells suppress mutagenic end joining is not well understood.

SPO11 forms DSBs in a highly regulated fashion early in meiotic prophase I^5^. In many species, SPO11 preferentially cuts DNA within narrow genomic segments called hotspots^6,7^. There are ∼15,000 hotspots in wild-type mice, but each meiocyte makes ∼250–350 DSBs, so only a small fraction of hotspots is used in any given meiosis^8-10^.

SPO11 remains covalently attached to the 5’ DNA ends after DNA cleavage. In yeasts, plants and mice, SPO11 is released still bound to a short oligonucleotide (oligo) by an endonucleolytic nick on the 5’ strand that is dependent, where tested, on MRE1^11-18^. Most SPO11 oligos in mice range ∼15–35 nt in length and have been sequenced to obtain nucleotide-resolution DSB maps^8,11,19^. NBS1, which together with MRE11 and RAD50 forms the MRN complex^20^, has also been implicated in SPO11 removal in mice^21^. Following SPO11-oligo removal, resection produces long 3’ single-stranded DNA (ssDNA) overhangs (mean ∼1100 bp in mice^22,23^) onto which RAD51 and DMC1 bind to catalyze strand invasion into homologous duplexes, forming early recombination intermediates^24^.

Central to DSB regulation is the conserved ATM kinase, which responds to DSBs by inhibiting further SPO11 activity both locally and genome-wide^19,25-29^. The ∼10-fold increase in meiotic DSBs with ATM loss in mouse spermatocytes is accompanied by much more frequent use of end joining, giving rise to deletions and tandem duplications at double cuts formed at adjacent hotspots, and to microdeletions at double cuts within single hotspots^4^. SPO11 double cutting within individual hotspots also liberates DNA fragments that can be inserted ectopically during end joining at distant hotspots^4^. The *de novo* indels and structural variants (SVs) that arise from double cutting are thus expected to contribute to genome variation^30,31^.

How SPO11-generated DSBs result in end joining is unclear. In *Caenorhabditis elegans* and *Drosophila melanogaster*, end joining resulting in chromosome aggregates becomes active in mutants with compromised DSB processing or recombination^32-37^, but whether such defects contribute to the mutagenic end joining we observe at double cuts in mammalian meiosis is unknown.

In addition to an increase in DSB numbers, *Atm*^−*/*–^ spermatocytes also show resection defects, displaying an increased frequency of both unusually long and unusually short resection tracts^22,23^. Additionally, a subpopulation of DSB ends remains unresected, attributed to the presumed role of ATM in regulating MRE11 endonuclease activity^22,23,38^. The complex dysregulation of both formation and processing of DSBs makes it difficult to determine which type of DSB ends are able to undergo deletions and other mutational outcomes by end joining.

To address the impact of SPO11-DSB processing, we have now mapped end-joining events initiated by double cutting in conditional *Mre11* knockout mice and hypomorphic MRN mutants. Our findings demonstrate that end joining can occur at unresected DSB ends, that MRE11-dependent SPO11 removal is not required for end ligation, and that an alternative mechanism for SPO11 removal influences deletional and insertional end joining. In addition, microdeletion and ectopic insertion data suggest that MRE11 and ATM collaborate to control the spatial organization of DSBs in and around hotspots. This study advances our understanding of DSB regulation and germline mutagenesis in mammalian meiosis.

## Results

### End joining at distantly spaced DSBs

Meiotic DSB hotspots are both numerous and unevenly spaced, creating opportunities for double cutting at adjacent hotspots separated by a wide range of distances. We previously demonstrated deletions from double cuts at hotspots separated by <2 kb^4^. These deletions included events that altered gene structure by eliminating an exon, so we reasoned that double cutting may be even more mutagenic if end joining can also occur between more distantly spaced DSBs. Relevant to this, 50% of autosomal hotspots defined in *Atm*^−*/*–^ SPO11-oligo maps^8^ are located within 30 kb of another hotspot^4^.

To test for larger deletions, we selected a DSB hotspot cluster on Chr13, present in both wild-type and *Atm*^−*/*–^ SPO11-oligo maps, which contains four hotspots of similar strength separated by various distances (**Supplemental Fig. 1A**). We performed nested PCR on testis DNA^4^ to detect deletions between these more distantly spaced hotspots. For each assayed hotspot pair—with distances of 5 kb, 15 kb, 20 kb, or 30 kb—PCR does not amplify parental DNA due to the short extension time (**Fig. 1A, B; Supplemental Fig. 1; Supplemental Table S1**). Genomic DNA equivalent to ∼16,000 haploid genomes from either ATM-proficient (*Atm*^*+/+*^ or *Atm*^*+/*–^) or -deficient testes was seeded in individual wells of 96-well plates to analyze a total of ∼4.8 million genome equivalents per hotspot pair (**Fig. 1B; Supplemental Table S2**). Deletion products were identified by gel electrophoresis and then sequenced to map deletion breakpoints.

**Figure 1.**
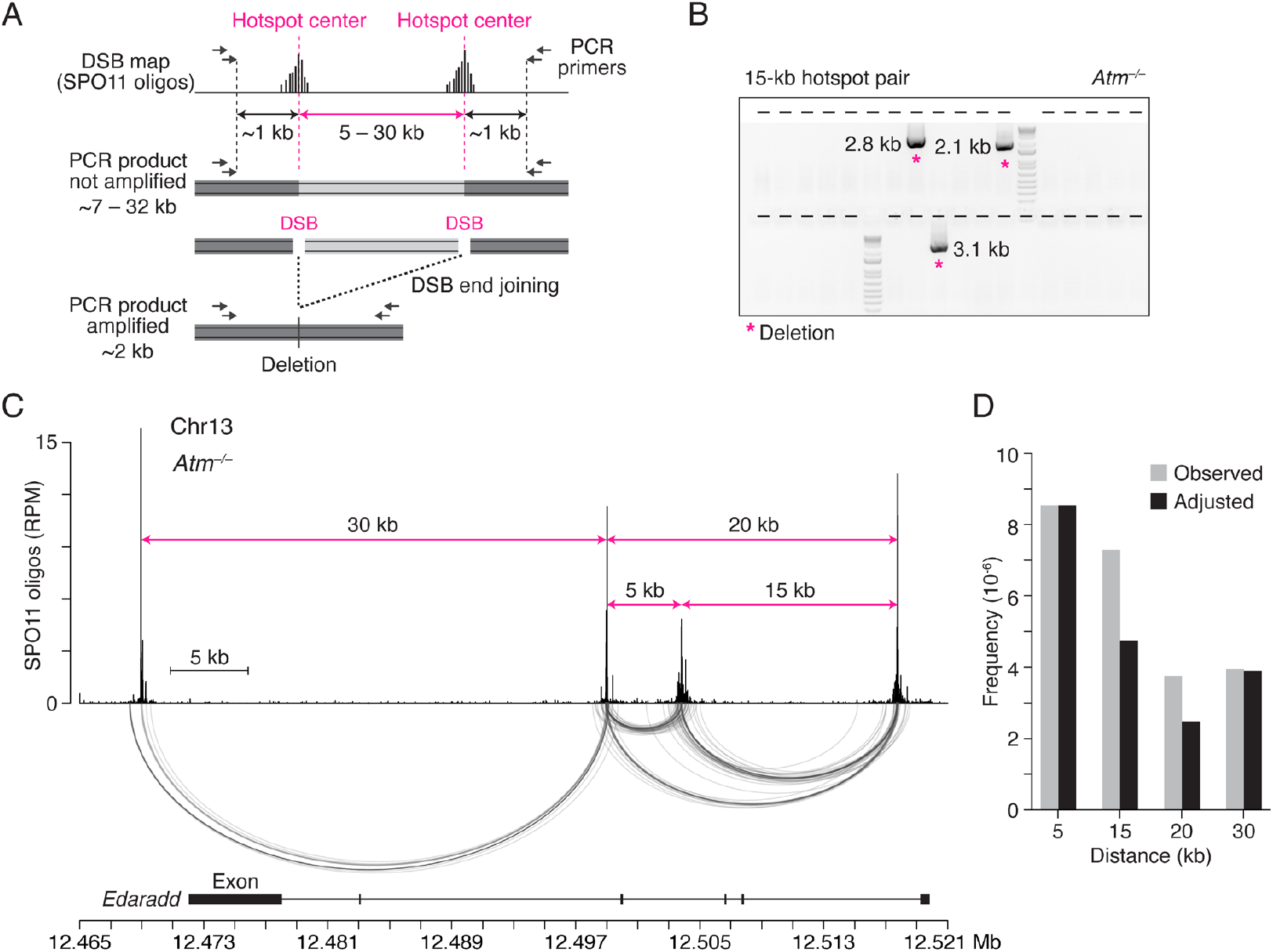
Double cuts at distantly spaced hotspots are efficiently joined in ATM-deficient spermatocytes. **A**. General strategy to detect deletions between meiotic DSB hotspots. Top: Schematic of SPO11-oligo maps at hotspot pairs separated by various distances. Dashed pink lines indicate hotspot centers. Bottom: Nested PCR assay to detect deletions. See also **Supplemental Fig. S1A**. Joining of DSB ends deletes the DNA between hotspots. **B**. representative agarose gel detecting deletions (asterisks) at the 15-kb hotspot pair in *Atm*^−*/*–^ spermatocytes. Each lane (indicated by tick marks) contains material from the secondary, nested PCR after a primary PCR seeded with ∼16,000 haploid genome equivalents. Most of the lanes lack an amplification product. PCR product sizes were determined by sequencing. See also **Supplemental Fig. S1B**. **C**. Similar distributions in *Atm*^−*/*–^ of deletion breakpoints and SPO11 oligos for the hotspot cluster on Chr13 (RPM, reads per million). SPO11-oligo maps are from *Atm*^−*/*–^ and are used throughout^8^. The distances between the tested hotspot pairs are demarcated by pink arrows. Arc diagrams below the SPO11-oligo map connect the breakpoints of individual deletions. Genomic coordinates (mm10) are shown below with the *Edaradd* exons. Each deletion removes one or more exons. **D**. Deletion frequencies in *Atm*^−*/*–^ spermatocytes at hotspots separated by increasing distances. Five to six mice were analyzed for each hotspot pair. Observed frequencies per haploid genome (gray) were adjusted (black) for hotspot strength (**Supplemental Fig. S1A**) by normalizing the expected double-cutting frequencies at the 15-, 20-, and 30-kb hotspot pairs to that of the 5-kb hotspot pair.

In *Atm*^−*/*–^ spermatocytes, deletions were detected at all tested hotspot pairs, including those separated by 20 kb and 30 kb (frequencies ranging from ∼4 to 9 × 10^−6^ per haploid genome) (**Fig. 1C; Supplemental Table S2**). The distribution of deletion breakpoints was similar to the distribution of SPO11 oligos, as expected from end joining at double cuts^4^. In ATM-proficient cells, only one deletion was observed (≤0.2 × 10^−6^) (**Supplemental Table S2**), indicating that ATM suppresses deletions at both closely and distantly spaced hotspots. The frequencies of deletions in *Atm*^−/–^ were similar for the 15-kb, 20-kb and 30-kb hotspot pairs when adjusted for hotspot strength, and were 1.8-to 3.5-fold lower than for deletions for the 5-kb hotspot pair (**Fig. 1D; Supplemental Table S2**). Thus, end joining can occur between distantly spaced DSBs. The lower deletion frequency for more distantly spaced DSBs may indicate that the efficiency of deletional end joining and/or the frequency of double cutting scales inversely with distance.

### End joining at unresected DSB ends in the absence of MRE11

To determine whether end joining can occur at unresected DNA ends, we tested for deletions in the absence of MRE11, which has a conserved role in DSB processing^11,15-18,32,34,39,40^. We used *Mre11* conditional knockout mice, in which a floxed *Mre11* allele^41^ is deleted in differentiating spermatogonia using *Ngn3-Cre*^42,43^. For simplicity, *Mre11*^−*/*–^ will refer to the *Mre11*^*flox/*–^ *Ngn3-Cre+* genotype, and *Mre11*^*+/*–^ will refer to the *Mre11*^*flox/*–^ or *Mre11*^*flox/+*^ *Ngn3-Cre+* genotypes. MRE11 deficiency results in spermatogenic failure and male infertility with the accumulation of unresected DSBs^18^. As expected, the lack of resection in *Mre11*^−*/*–^ results in highly similar *Mre11*^−*/*–^ S1-seq and SPO11-oligo average profiles^18^, although at individual hotspots, peak positions and heights do not exactly match, including differences between strand-specific S1-seq signals (**Supplemental Fig. 2A**).

Deletions were readily detected at both the 5-kb and 15-kb hotspot pairs in *Mre11*^−*/*–^ spermatocytes but were rare in control *Mre11*^*+/*–^ cells (**Fig. 2A; Supplemental Table S3**). Frequencies in *Mre11*^−*/*–^ were similar to those in *Atm*^−*/*–^, consistent with MRE11 regulating DSB numbers^18,44-46^, likely through its role in activating ATM^47,48^. Strikingly, deletion break-points in *Mre11*^−*/*–^ showed distinctive distributions compared to *Atm*^−*/*–^, with breakpoints more tightly clustered around hotspot centers (**Fig. 2A**), which was particularly evident in cumulative distribution plots (**Fig. 2B**). Within 200 bp around the hotspot centers, the fraction of breakpoints was higher in *Mre11*^−*/*–^ (75–88%) than in *Atm*^−*/*–^ (34–65%). Furthermore, within these regions, breakpoint peaks in *Mre11*^−*/*–^ and *Atm*^−*/*–^ overlapped and matched SPO11-oligo peaks, but the *Mre11*^−*/*–^ peaks were more prominent (**Fig. 2C, Supplemental Fig. 2B**). These results suggest that unresected DSB ends can undergo end joining in *Mre11*^−*/*–^ cells.

**Figure 2.**
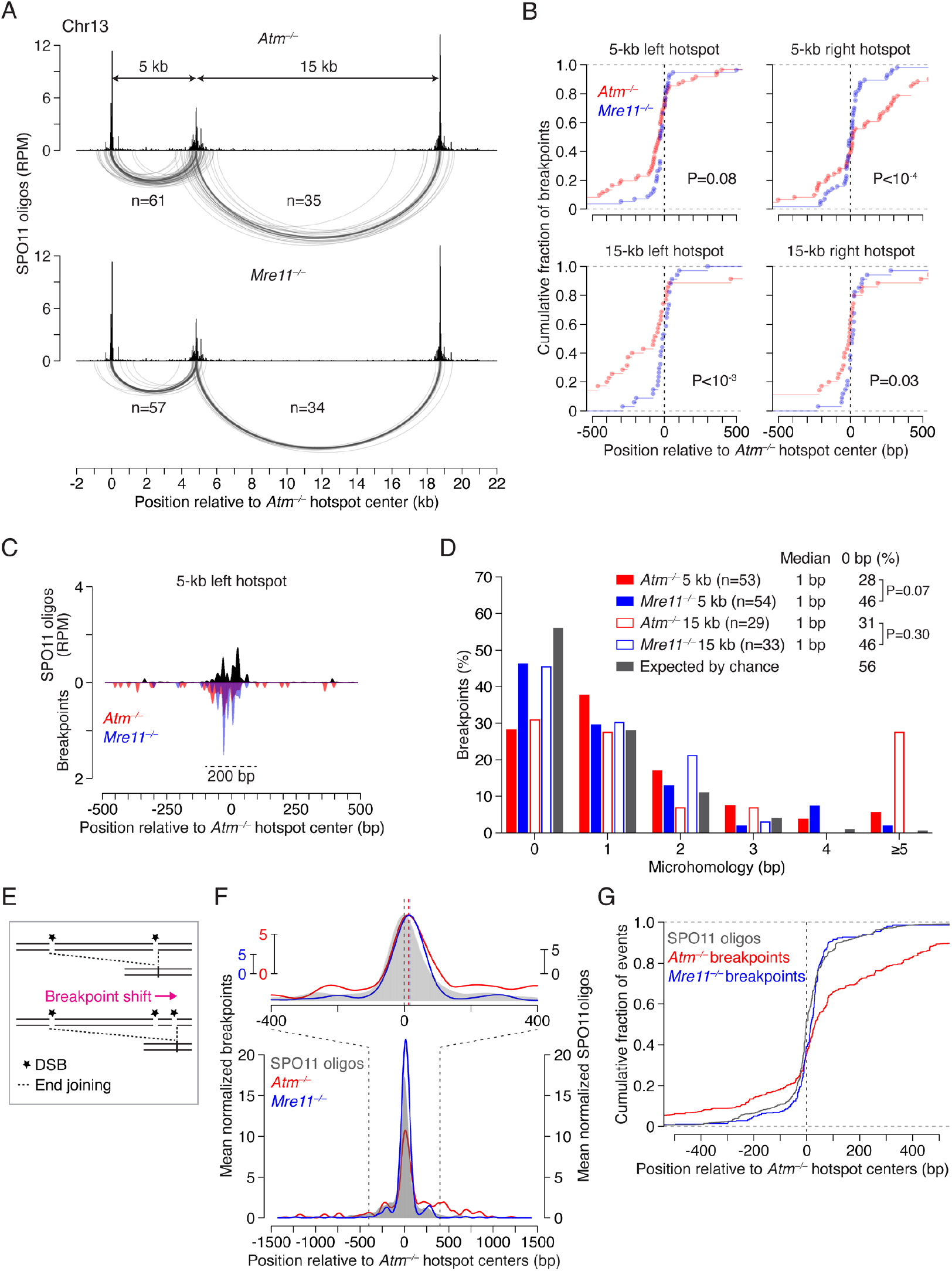
Deletions arise at unresected double cuts in the absence of MRE11. **A**. Arc diagrams of deletions in *Atm*–*/*– and *Mre11*–*/*– at the 5-kb and 15-kb hotspot pairs. A similar number of deletions were analyzed per genotype. **B**. Cumulative distributions of breakpoints in *Atm*–*/*– and *Mre11*–*/*– at individual hotspots. P values are from **Levene’s test**. **C**. Smoothed breakpoint and SPO11-oligo profiles (21-bp Hann filter). Breakpoints are normalized to the number of deletions in *Atm*–*/*–. Dashed line indicates a 200-bp window around the hotspot center. For other hotspots see **Supplemental Fig. 2B**. **D**. Distribution of microhomology lengths. Microhomology expected by chance was calculated as previously described70. Deletions with insertions were excluded. P values are from **Fisher’s exact test**. **E**. Expected impact of double cutting within hotspots on end-joining breakpoint distributions. In this example, double cutting within the right hotspot shifts the deletion breakpoint to the right. **F**. Mean normalized breakpoint and SPO11-oligo profiles across five hotspots in three pairs: the 5-kb and 15-kb hotspot pairs on Chr13, and a 2-kb hotspot pair on Chr1 (**Supplemental Fig. 3A**). Data were locally normalized by dividing the signal at each base pair by the mean signal within a 3001-bp window for each hotspot, and then averaged across all hotspots. Signals for the left hotspot in each pair were flipped prior to averaging. The top panel shows a zoom into the region around hotspot centers, with peak heights matched and peak positions marked with vertical dashed lines. Profiles are smoothed with a 151-bp Hann filter. **G**. Cumulative distributions of breakpoints and SPO11 oligos (data from panel **F**).

Joining of unresected ends in the absence of MRE11 would be expected to lead to fewer microhomologies at deletion breakpoints due to the presence of 2-nt 5’ overhangs generated by SPO11 cleavage, in contrast to long 3’ ssDNA overhangs that could expose such sequences in *Atm*^−*/*–^. While microhomologies were generally short in both *Atm*^−*/*–^ and *Mre11*^−*/*–^ (median of 1 bp), breakpoints with 0 bp microhomology at both the 5-kb and 15-kb hotspot pairs tended to be more frequent in *Mre11*^−*/*–^ (∼46%) than in *Atm*^−*/*–^ (∼26–31%), and were only slightly lower than the 56% expected by chance (**Fig. 2D**). In addition, for the 15-kb hotspot pair, we observed frequent deletions with longer microhomologies (≥5 bp) in *Atm*^−*/*–^ (28%; 8/29), but not in *Mre11*^−*/*–^ (0%; 0/33; P = 0.0045, Fisher’s exact test). Of those eight deletions in *Atm*^−*/*–^, two involved recurrent use of a 10-bp microhomology, similar to what we observed at other hotspot pairs previously^4^. These observations are thus consistent with joining of unresected ends in the *Mre11* mutant—likely through nonhomologous end-joining (NHEJ)—and joining of some resected ends in the *Atm* mutant, possibly through microhomology-mediated end joining (MMEJ)^2^.

### End-joining breakpoint distributions are shaped by double cutting within hotspots

In addition to double cutting at adjacent hotspots, double cuts can also form within individual hotspots^4,18,25,49^, and our previous findings in *Atm*^−*/*–^ mice suggested that their occurrence shapes deletion breakpoint distributions at hotspot pairs^4^. We reasoned that the impact of double cutting on end-joining profiles would be apparent in the absence of MRE11 due to the lack of resection. Double cutting within a hotspot should cause end-joining breakpoints to shift outward from the hotspot center, in the direction away from the other hotspot in the pair (**Fig. 2E**).

To be able to observe subtle shifts in breakpoint distributions from double cutting, we computed average profiles of breakpoints and SPO11 oligos using three hotspot pairs (**Fig. 2F**). A clear pattern emerged in *Mre11*^−*/*–^ in that the major breakpoint peak was shifted to the right of the SPO11-oligo peak. In cumulative distribution plots, this shift had a median of 18 bp (P = 2.89 × 10^−10^, Wilcoxon test) (**Fig. 2G**). In addition, the secondary right breakpoint peak was stronger than the left secondary peak. These results support the prediction that breakpoint distributions are influenced by double cutting within single hotspots in the absence of MRE11 (see also below).

In *Atm*^−*/*–^, a shift of the major breakpoint peak was also observed (**Fig. 2F**). Overall, breakpoints were shifted to the right by a median of 33 bp (P = 2.2 x 10^−16^, Wilcoxon test) (**Fig. 2G**), again implicating a contribution of double cutting. However, many breakpoints were spread away from the hotspot center, a pattern also seen at individual hotspots^4^ (**Supplemental Fig. 2B,C; 3A**), unlike in *Mre11*^−*/*–^ where the pattern was more centered. Considering the double-cutting signature in *Mre11*^−*/*–^, we infer that the wider distribution of breakpoints in *Atm*^−*/*–^ (**Fig. 2F,G**) largely reflects joining of resected ends (spread to the right) and, likely, resected ends extended by DNA synthesis after strand invasion (spread to the left).

### TDP2 removal of SPO11 in the absence of MRE11

For blocked DSB ends to be joined in *Mre11*^−*/*–^ spermatocytes, SPO11 must be removed through another mechanism. We considered the tyrosyl-DNA phosphodiesterase TDP2 because of its role in cleaving 5’ DNA covalent linkages of TOP2 (topoisomerase II) in mitotic cells^50,51^. Mammalian TDP2 can remove SPO11 from DNA ends *in vitro*, including in *Atm*^−*/*–^ and *Mre11*^−*/*–^ testis DNA samples^22,25,49,52,53^.

To test whether TDP2 can foster end ligation *in vivo*, we examined *Mre11*^−*/*–^ *Tdp2*^−*/*–^ double mutant mice for deletions at the 5-kb hotspot pair and the previously studied 2-kb hotspot pair on Chr1^4^. In *Mre11*^−*/*–^ single mutants, deletions at the 2-kb hotspot pair were observed at a similar frequency as in the absence of ATM and with narrower breakpoint profiles (**Fig. 3A; Supplemental Fig. 3A; Supplemental Table S4**), similar to the deletions within the Chr13 hotspot cluster. In *Mre11*^−*/*–^ *Tdp2*^−*/*–^ double mutants, deletions were detected at both the 2-kb and 5-kb hotspot pairs (**Fig. 3A; Supplemental Fig. 3B**), but deletion frequencies were reduced ∼2-fold relative to *Mre11* single mutants (**Supplemental Tables S3 and S4**). These results indicate that TDP2 promotes end joining, presumably by removing SPO11 from DSB ends, but that an alternative mechanism(s) must also exist. We further infer that both TDP2-dependent and -independent SPO11 removal from DSB ends in the *Mre11*^−*/*–^ background occurs without substantial loss of DNA end sequences, because deletion breakpoint profiles overlap in the double and single mutants (**Fig. 3A,B; Supplemental Fig. 3B,C**).

**Figure 3.**
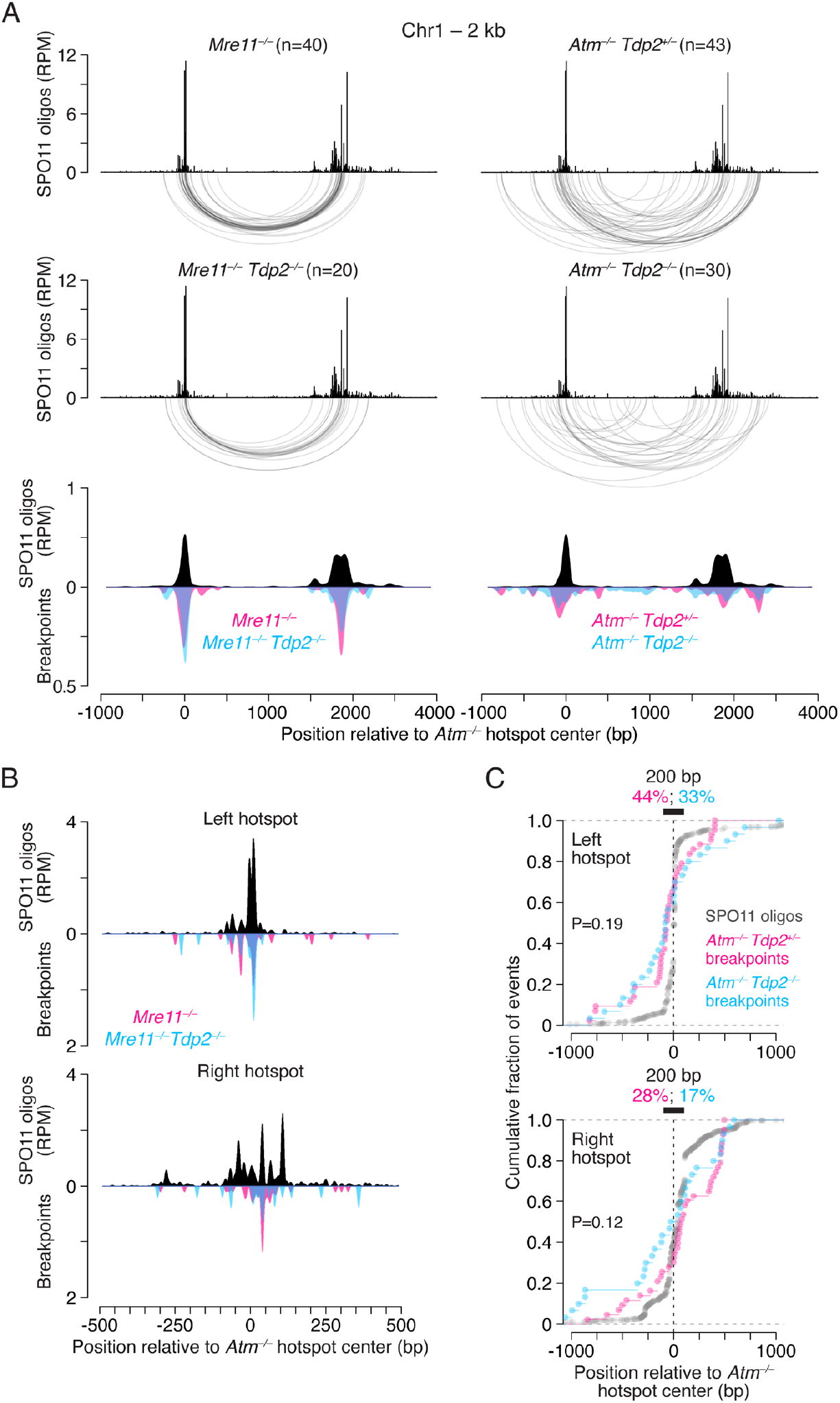
TDP2 promotes deletions in the absence of MRE11-mediated SPO11 removal. **A**. Comparison of deletion breakpoint distributions in *Mre11*^−*/*–^ and *Atm*^−*/*–^ single mutants with *Mre11*^−*/*–^ *Tdp2*^−*/*–^ and *Atm*^−*/*–^ *Tdp2*^−*/*–^ double mutants at the 2-kb hotspot pair on Chr1. Below are the corresponding smoothed breakpoint and SPO11-oligo profiles (151-bp Hann filter). Breakpoints are normalized to the number of deletions in *Atm*^−*/*–^ *Tdp2*^*+/*–^. **B**. Smoothed breakpoint and SPO11-oligo profiles close to hotspot centers (21-bp Hann filter) for the 2-kb hotspot pair on Chr1. Breakpoints are normalized to the number of events in *Mre11*^−*/*–^. **C**. Cumulative distributions of breakpoints and SPO11 oligos at the 2-kb hotspot pair on Chr1. For SPO11 oligos, reads mapping within ±1500 bp from hotspot centers were included. Black bars indicate a 200-bp window around the hotspot centers, with the percentage of breakpoints within this region shown above. P values are from **Levene’s test**. Similar results were obtained for the 5-kb hotspot pair on Chr13 (see **Supplemental Fig. 3B–D**).

We also asked whether TDP2 promotes the end joining that occurs in *Atm*^−*/*–^. However, because only a fraction of DSBs remain unresected (SPO11-blocked) in the absence of ATM, loss of TDP2 is expected to have less of an impact compared to loss of MRE11. Deletion frequencies showed high variability between mice but did not show a consistent reduction in frequency (**Supplemental Table S5**). In *Atm*^−*/*–^ *Tdp2*^−*/*–^ double mutants, breakpoint profiles at both hotspot pairs were slightly more spread out compared to *Atm*^−*/*–^ *Tdp2*^*+/*–^ (**Fig. 3A; Supplemental Fig. 3B**), such that the percentage of breakpoints within 200 bp around hotspots centers decreased from ∼28–44% in *Atm*^−*/*–^ *Tdp2*^*+/*–^ to ∼17–36% in *Atm*^−*/*–^ *Tdp2*^−*/*–^ (**Fig. 3C; Supplemental Fig. 3D**). Therefore, it is plausible that TDP2 contributes to SPO11 removal at some DSBs in the *Atm*^−*/*–^ mutant.

### MRE11 is dispensable for ectopic insertions of double-cut fragments

In the absence of ATM, a subset of deletions also had insertions of two distinct types: an ectopic fragment from another hotspot, likely originating from a double cut, or a locally inverted sequence^4^ (**Fig. 4A)**. The inverted insertions may arise from resolution of foldback structures or from reinsertion of double-cut fragments in inverted orientation. Unlike ectopic insertions, each inverted insertion displayed microhomology at one of its breakpoints, implying a requirement for end resection to reveal that microhomology^4^. We therefore anticipated that local inversions, but not ectopic insertions, would be reduced or absent when only unresected DSBs are present.

**Figure 4.**
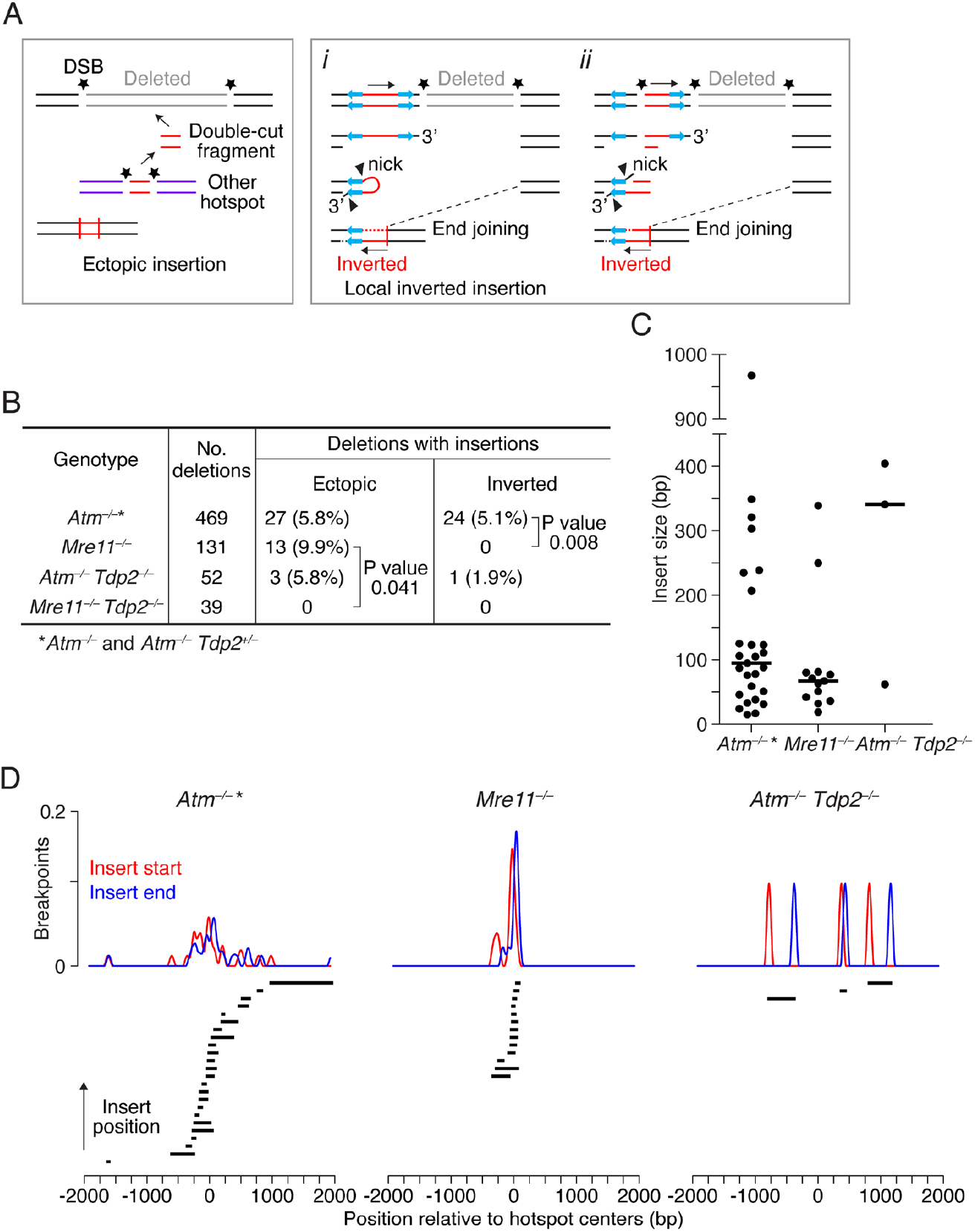
Ectopic insertions of double-cut fragments occur in the absence of MRE11 and require TDP2. **A**. Potential origins of insertions. Left, ectopic insertion of a fragment generated by double cutting at another hotspot. Right, local inverted insertion involving either foldback of a resected DSB end (*i*) or a triple cut (*ii*), promoted by the presence of short inverted sequences. **B**. Frequency of deletions with ectopic or inverted insertions across genotypes. Insertions in *Atm*^−*/*–^ include previously published data^4^. Ectopic inserts that did not map to other hotspots or could not be mapped were excluded (4 in *Atm*^−*/*–^, 1 in *Mre11*^−*/*–^ *Tdp2*^−*/*–^). P values are from Fisher’s exact test. **C**. Length distribution for ectopic insertions. Median insert lengths are indicated. *Atm*^−*/*–^ * inserts are from both *Atm*^−*/*–^ *Tdp2*^*+/*–^ and *Atm*^−*/*–^, which includes some that were previously mapped^4^. **D**. Positions of inserts relative to their source hotspot centers. Smoothed breakpoint profiles (151-bp Hann filter) are normalized to the number of events in *Atm*^−*/*–^.

Consistent with prediction, all of the insertions in *Mre11*^−*/*–^ mice (9.9% of all deletion events, 13/131) were ectopic, and none derived from locally inverted sequences (**Fig. 4B**). Ectopic insertions in *Mre11*^−*/*–^ were somewhat more frequent than in *Atm*^−*/*–^ (5.8%), but this difference was not significant (P = 0.111, Fisher’s exact test). However, the absence of inverted insertions in *Mre11*^−*/*–^ was significantly different from *Atm*^−*/*–^, for which 5.1% of deletions had inverted insertions (24/469, P = 0.004. This finding supports the conclusion that end resection is required for inverted insertions.

The ectopically inserted DNA sequences in *Mre11*^−*/*–^, similar to *Atm*^−*/*–^, derived from genomic sites showing SPO11 activity, either to SPO11-oligo hotspots (12/13) or to a weaker SPO11-oligo cluster that did not meet the hotspot-definition threshold (1/13), so we collectively classified all 13 as originating from other hotspots. Finding inserts in the absence of MRE11 supports the premise that they represent double-cut fragments and excludes an alternative DNA synthesis-based insertion mechanism (**Supplemental Fig. 4A**), which would require resection for strand annealing. We conclude that double-cut fragments do not require MRE11-mediated processing prior to insertion.

Inserts from other hotspots were generally small in both *Atm*^−*/*–^ (15– 967 bp; median 95 bp) and *Mre11*^−*/*–^ (19–339 bp; median 67 bp) (**Fig. 4C**). Given that in yeast Mre11 does not cleave Spo11 from double-cut fragments <70 bp^49^, these small SPO11-capped fragments may be stabilized, enhancing their potential for mutagenic insertions in mice.

Remarkably, unlike in *Mre11*^−*/*–^ single mutants, we did not capture ectopic insertions from other hotspots in *Mre11*^−*/*–^ *Tdp2*^−*/*–^ double mutants (P = 0.041) (**Fig. 4B**). Thus, TDP2 may be required to cleave SPO11 from small double-cut fragments to promote their insertion. Thus, this contrasts with the occurrence of deletions in *Mre11*^−*/*–^ *Tdp2*^−*/*–^ double mutants (**Fig. 3A**), where SPO11-capped ends can be processed by another mechanism(s). Interestingly, in the *Atm*^−*/*–^ *Tdp2*^−*/*–^ double mutants, 2 of the 3 ectopic insertions were relatively large at 341 bp and 404 bp (**Fig. 4C**), and thus may have utilized MRE11 for SPO11 removal.

When we further analyzed how ectopically inserted sequences were positioned at the hotspots from which they originated, *Mre11*^−*/*–^ inserts mapped much closer to hotspot centers than *Atm*^−*/*–^ inserts (**Fig. 4D**). The wider distribution in *Atm*^−*/*–^ was not due to the higher proportion of larger inserts, as even smaller *Atm*^−*/*–^ inserts were found at more variable distances from hotspot centers (**Supplemental Fig. 4B**).

### MRE11 suppresses clustered microdeletions within hotspots

We envisioned that the narrower distribution of source positions for the ectopic inserts in the absence of MRE11 results from more tightly clustered double cuts, which could also lead to small intra-hotspot deletions, termed microdeletions. To test this, we applied an amplicon deep sequencing strategy we previously used to detect microdeletions in the absence of ATM^4^. Four overlapping amplicons of ∼580 bp each spanning ∼900 bp of a strong hotspot on Chr1 were amplified from testis DNA equivalent to ∼10 million haploid genomes per amplicon. Multiple PCRs were pooled for each amplicon and sequenced to identify reads containing microdeletions and to map breakpoints.

Collectively across amplicons, we identified 3,317 microdeletion reads in *Mre11*^−*/*–^ (158.6 microdeletions per million reads) and 647 microdeletion reads in earlier *Atm*^−*/*–^ data^4^ reanalyzed with the same pipeline (54.9 microdeletions per million reads) (**Supplemental Table S8, Dataset S1; Methods**). Considering that identical reads may represent PCR duplicates of the same input DNA molecule, we conservatively filtered unique microdeletions for each amplicon and identified 804 and 309 independent microdeletions in *Mre11*^−*/*–^ and *Atm*^−*/*–^, respectively (**Supplemental Table S8; Dataset S1**). Microdeletion lengths had similar ranges in *Mre11*^−*/*–^ (11–339 bp) and *Atm*^−*/*–^ (11–361 bp) (**Fig. 5A; Supplemental Fig. 5A, 6A**). Thus, microdeletions are abundant in the absence of either MRE11 or ATM.

**Figure 5.**
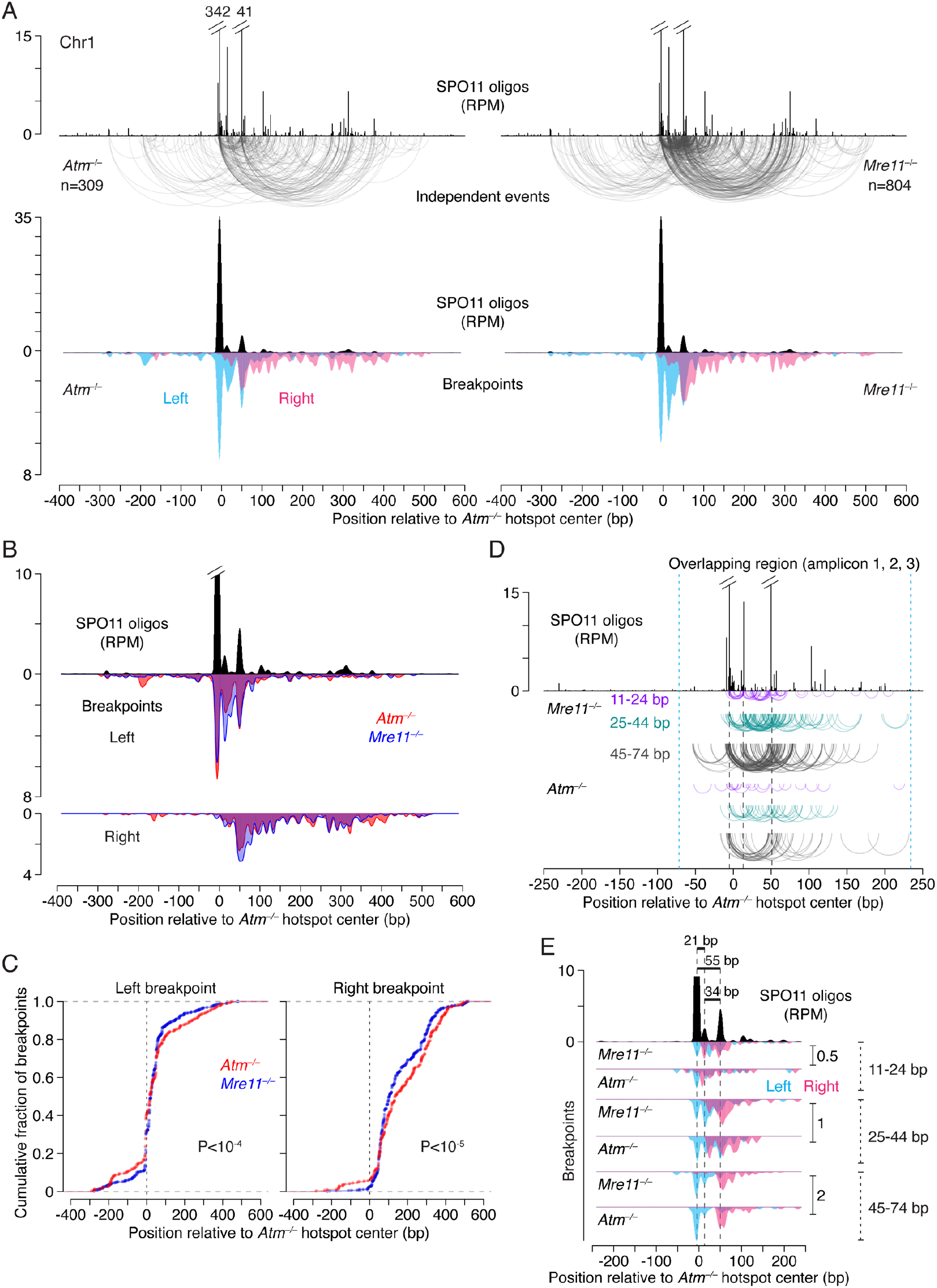
Microdeletions cluster within a single hotspot in the absence of MRE11. **A** Microdeletions at small double cuts detected in *Mre11*^−*/*–^ and *Atm*^−*/*–^ using amplicon deep sequencing. Top: SPO11-oligo maps and arc diagrams of independent microdeletions identified in four amplicons. The SPO11-oligo RPM values for the two highest peaks are indicated. Bottom: Smoothed microdeletion breakpoint and SPO11-oligo profiles (21-bp Hann filter). Breakpoints are normalized to the number of events in *Atm*^−*/*–^. *Atm*^−*/*–^ microdeletions are from reanalyzed published data^4^. See **Supplemental Fig. 5A,B** and **Supplemental Table S8** for microdeletion distributions in each amplicon and for microdeletion frequencies, respectively. **B** Comparison of smoothed left and right breakpoint maps between *Mre11*^−*/*–^ and *Atm*^−*/*–^, relative to SPO11-oligo maps (21-bp Hann filter). Breakpoints are normalized to *Atm*^−*/*–^ events. **C** Cumulative distributions of left and right breakpoints. P values are from **Levene’s test**. **D** Distributions of microdeletions (arc diagrams) stratified by length in the overlapping region of amplicons 1, 2, and 3. Dashed vertical blue lines delineate the 304-bp overlapping region between the forward primer of amplicon 3 and the reverse primer of amplicon 1 (**Supplemental Fig. 5A**). Dashed black lines are at the major and minor SPO11-oligo peaks, as in panel **E**. **E** Comparison of the SPO11-oligo profile with the breakpoint distributions of microdeletions stratified by length. Distances between the major and minor SPO11-oligo peaks (dashed lines) are indicated. Breakpoint profiles are normalized to the number of microdeletions in *Atm*^−*/*–^. Maps are smoothed with a 21-bp Hann filter.

Microdeletion breakpoint profiles for *Mre11*^−*/*–^ and the reanalyzed *Atm*^−*/*–^ data reflected SPO11-oligo profiles, with the major left and right breakpoint peaks overlapping the major and secondary SPO11-oligo peaks, respectively (**Fig. 5A**), consistent with double cutting as the initiating event for microdeletions in both mutants. *Mre11*^−*/*–^ and *Atm*^−*/*–^ end-joining profiles showed broad overlap between left or right break-point peak positions (**Fig. 5B**), which is even more evident in the profiles with combined left and right breakpoints (**Supplemental Fig. 5C**). Such overlap would not be expected if joining occurred at variably resected DSB ends, suggesting that in the absence of either MRE11 or ATM, most microdeletions form at unresected or very minimally processed DSB ends. Supporting this notion, even minor breakpoint peaks frequently overlapped SPO11-oligo peaks (**Fig. 5B; Supplemental Fig. 5B**).

Nonetheless, we observed subtle differences in the breakpoint distributions between the two mutants. Specifically, *Mre11*^−*/*–^ showed more prominent signal both for the left breakpoint peaks between the major and secondary SPO11-oligo peaks and for the major right breakpoint peak, while *Atm*^−*/*–^ breakpoint profiles were modestly, but highly significantly, more spread out (**Fig. 5B, C**). These differences in breakpoint distributions were evident for individual amplicons as well (**Supplemental Fig. 5B**). Of note, a higher fraction of microdeletions in *Mre11*^−*/*–^ (29%; 234/804) than in *Atm*^−*/*–^ (21%; 66/309; P = 0.0102, Fisher’s exact test) had both breakpoints mapping within the 100 bp segment encompassing the major SPO11-oligo peaks (from −25 bp to +75 bp relative to the hotspot center). Thus, microdeletions cluster more tightly around the hotspot center in the absence of MRE11 than in the absence of ATM. Importantly, since the more spread-out breakpoint peaks in *Atm*^−*/*–^ frequently overlapped weaker *Mre11*^−*/*–^ peaks (unresected DSB ends), this suggests that double cuts themselves cluster more tightly in *Mre11*^−*/*–^. However, we cannot entirely exclude that, as proposed for deletions between hotspots, the slightly wider *Atm*^−*/*–^ profiles reflect joining at resected ends or at extended strand-invading ends.

As previously observed in *Atm*^−*/*–4^, density plots of microdeletion sizes in *Mre11*^−*/*–^ showed prominent peaks at ∼30 bp and ∼60 bp comprising ∼30% and ∼40% of events in *Atm*^−*/*–^ and *Mre11*^−*/*–^, respectively (**Supplemental Fig. 6A**). These are particularly evident when analyzing events within the 304-bp region where amplicons 1, 2, and 3 overlap, as this region encompasses a majority of the DSBs at this hotspot (**Supplemental Fig. 6B**). The 30-bp microdeletion class is consistent with the minimum size of double cuts determined in prior studies: In both yeast and mice, the distribution of distances between the two DSBs in double cuts displays a ∼10-bp periodicity beginning at ∼3 helical turns, likely reflecting steric constraints that govern how close two co-oriented SPO11 complexes can be to one another^18,25,49,53-56^.

Surprisingly, however, we also observed a substantial number of microdeletions that were smaller than the expected minimum size, with events ranging from 11–24 bp accounting for ∼10% of microdeletions in both *Mre11*^−*/*–^ and the reanalyzed *Atm*^−*/*–^ data (**Supplemental Fig. 6B**). Very small microdeletions showing a peak at ∼15 bp in density plots were particularly apparent in *Atm*^−*/*–^, whereas in *Mre11*^−*/*–^, closer analysis revealed two distinct classes of ∼13 bp (11–17 bp) and ∼21 bp (18–24 bp) events (**Supplemental Fig. 6B,C**).

We further examined the spatial distribution of microdeletions grouped by length to bracket the <25 bp, ∼30 bp, and ∼60 bp size ranges. Similar to our previous study in *Atm*^−*/*–4^, in the absence of MRE11 the breakpoints of ∼30-bp and ∼60-bp microdeletions frequently clustered at prominent SPO11-oligo peaks that are separated by 34 bp and 55 bp, respectively (**Fig. 5D,E**). The <25 bp microdeletions clustered in the region between the major and secondary SPO11-oligo peaks in *Mre11*^−*/*–^ and to a lesser extent in *Atm*^−*/*–^ (**Fig. 5D,E**). In *Mre11*^−*/*–^, several breakpoints in the 18-24 bp class mapped precisely to the major SPO11-oligo peak and the adjacent peak 21 bp away (**Supplemental Fig. 6D**). This finding argues that double cuts separated by as little as two helical turns can occur in vivo and be joined. The class of exceptionally small microdeletions (<18 bp) in both *Mre11*^−*/*–^ and *Atm*^−*/*–^ have breakpoints that also cluster between the major and secondary SPO11-oligo peaks; possible sources of these microdeletions are addressed in the Discussion.

On the other extreme, the refined analysis now revealed frequent microdeletions of ≥200 bp in both mutants, most of which had one break-point near the hotspot center (**Supplemental Fig. 5A**). We also note that a fraction of ectopic inserts ranged from ∼200–350 bp (**Fig. 4C**), suggesting that when larger double cuts form within hotspots, they have the potential to reincorporate into the genome at other hotspots.

### Reducing resection length is not sufficient to provoke end joining

To further understand how DSB resection affects the propensity toward deletion formation, we asked whether end joining is impacted in mutants that initiate resection but have shortened resection tracts. MRE11 with the active site mutation H129N was expected to lack nuclease activity^57^, resulting in little or no resection initiation without affecting ATM signaling^41^ and hence meiotic DSB numbers. However, a parallel study found that only a small population of DSBs had unresected ends in *Mre11*^*H129N/flox*^ *Ngn3-Cre+* mice (simplified as *Mre11*^*H129N/*–^), with the large majority of DSBs instead showing a reduction in resection lengths by nearly half^18^. These results were likely caused by persistence of minimal amounts of wild-type MRE11 protein due to the use of a conditional allele necessitated by the embryonic lethality of *Mre11*^*H129N / H129N*^ mice^41^, although other interpretations are possible.

In *Mre11*^*H129N/*–^, the frequency of deletions at the 5-kb hotspot pair on Chr13 (0.71 × 10^−6^) was similar to that in *Mre11*^*+/*–^ (0.63 × 10^−6^) (**Supplemental Table S6**). The much lower deletion frequency compared to the absence of MRE11 is consistent with normal ATM activation and thus DSB levels^18^. These findings show that shorter resection lengths on their own are not sufficient to promote mutagenic end joining.

We further tested the impact of reduced resection using mice carrying a hypomorphic *Nbs1* allele, *Nbs1*^*ΔB* 58,59^. *Nbs1*^*ΔB/ΔB*^ spermatocytes show a resection profile similar to *Mre11*^*H129N/*– 18^ although with a small increase (1.8-fold) in SPO11 oligos^44^, consistent with reduced ATM activity^59^. Nonetheless, deletion frequencies were similar to *Mre11*^*H129N/*–^ and to controls at the 5-kb hotspot pair on Chr13, as well as at the 2-kb hotspot pair on Chr19^4^ (**Supplemental Tables S6, S7**).

Breakpoints in the few deletions obtained in *Mre11*^*H129N/*–^ and *Nbs1*^*ΔB/ΔB*^ were distributed broadly, similar to controls (**Fig. 6A,B**; **Supplemental Fig. 7A**). Collectively considering all events from mice that were neither *Atm*^−*/*–^ nor *Mre11*^−*/*–^ (32 combined events at the 5-kb hotspot pair and 27 at the 2-kb hotspot pair), deletion breakpoints did not cluster around hotspot centers but were more spread out than in *Atm*^−*/*–^ or *Mre11*^−*/*–^ (**Fig. 6C**; **Supplemental Fig. 7B**), as previously reported for a smaller number of events recovered at the 2-kb hotspot pair on Chr19^4^. Since we do not observe a broad overlap of SPO11-oligos and breakpoint peaks from controls and hypomorphs, joining may have occurred between resected ends, although we cannot exclude other scenarios.

**Figure 6.**
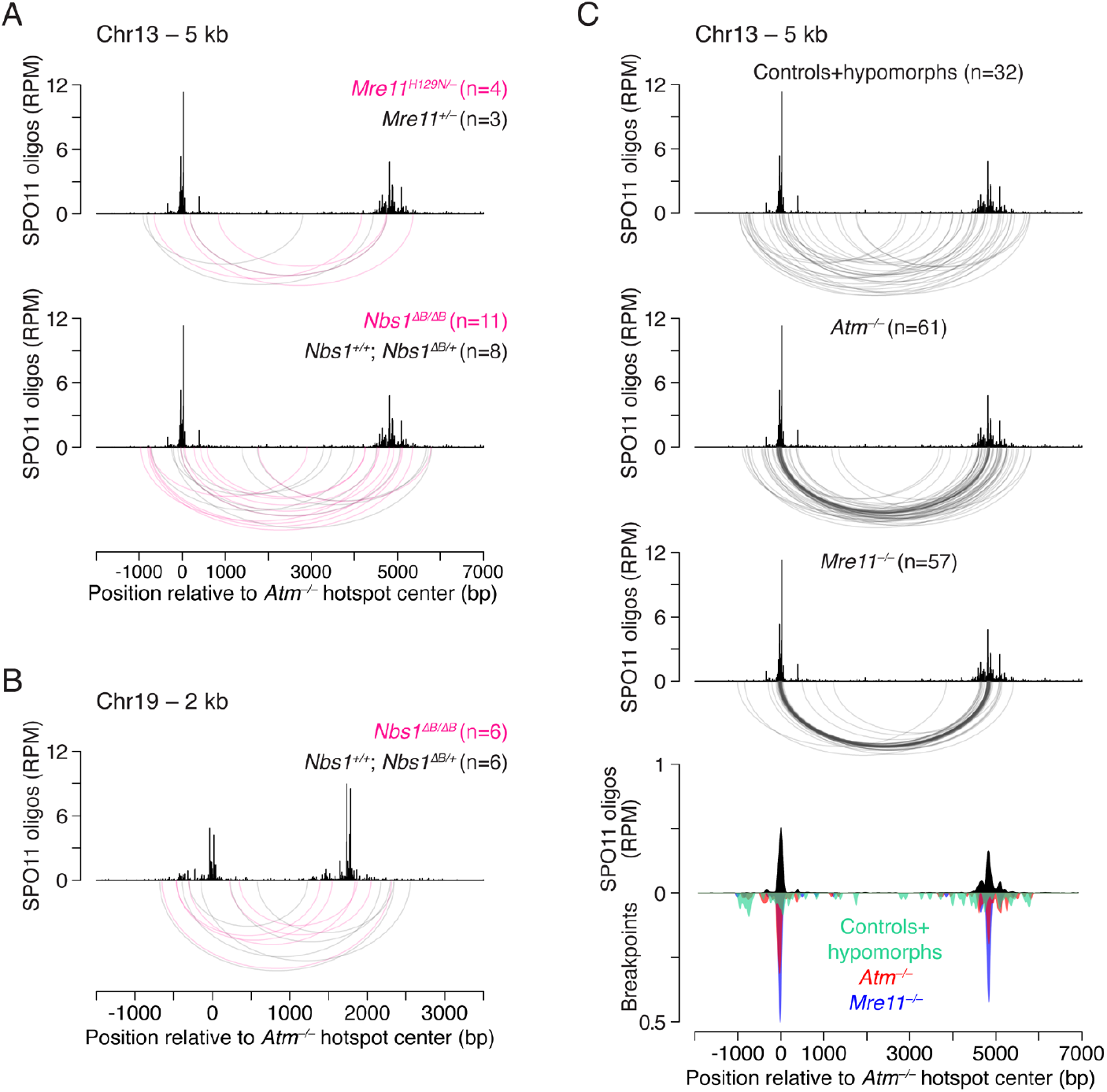
Deletion breakpoints in *Mre11*^*H129N/*–^ and *Nbs1*^*ΔB/ΔB*^, as in controls, do not cluster at hotspot centers. **A,B**. Deletion breakpoint distributions at the 5-kb hotspot pair on Chr13 (**A**) and the 2-kb hotspot pair on Chr19 (**B**) in the indicated genotypes (**Supplemental Tables S6, S7**). **C**. Comparison of *Atm*^−*/*–^ and *Mre11*^−*/*–^ breakpoint distributions to the combined distribution from controls (including from other experiments) and hypomorphic *Mre11*^*H129N/*–^ and *Nbs1*^*ΔB/ΔB*^ mice at the 5-kb hotspot pair. Breakpoints and SPO11-oligo maps are smoothed with a 151-bp Hann filter in the bottom graph. Breakpoints are normalized to the number of events in *Atm*^−*/*–^. Similar results were obtained for the 2-kb hotspot pair on Chr19 (see **Supplemental Fig. 7B**).

## Discussion

Our results uncover new features of meiotic end joining in mice. We show that deletions can arise at more distantly spaced hotspots, providing further evidence that double cutting is mutagenic and can encompass large genomic regions (at least 30 kb). We find that unresected DSBs can undergo end joining, such that in the absence of MRE11, breakpoint maps closely resemble SPO11-oligo maps. Further, TDP2 appears able to cleave SPO11 from DSB ends prior to ligation. This activity of TDP2 is not strictly required for DSB end joining but it appears to be essential to promote ectopic insertions of double-cut fragments. Finally, our findings demonstrate that MRE11 suppresses clustered microdeletions at double cuts within single hotspots, including at previously unanticipated SPO11 DSBs separated by as little as ∼21 bp.

### End joining at unresected double cuts

Loss of DSB control in the absence of ATM^8,19,26^ is associated with more abundant end joining events compared to wild type^4^. Prior experiments in *Atm*^−/–^ mice provided evidence that end joining can occur at resected DSBs but did not address whether unresected DSBs can also be substrates for joining. The studies of *Mre11*^−/–^ mice presented here show that unresected DSBs can indeed be joined. We therefore infer that *Atm*^−/–^ deletions are likely to arise at both resected and unresected DSBs. Further, we found in *Mre11*^−/–^ a footprint of small double cuts within single hotspots and determined the extent to which they shape deletion distributions at hotspot pairs, leading us to postulate that the wider deletion profiles in *Atm*^−/–^ result from joining of resected and extended strand-invading ends.

If we assume that DSBs and double cuts are similarly elevated in *Atm*^−/–^ and *Mre11*^−/–^ spermatocytes due to loss of MRE11-mediated activation of ATM^4,18,19,25^, it might imply that the propensity for end joining at double cuts is not significantly affected by the resection status of the DSBs. This inference arises because deletion frequencies at hotspot pairs are similarly increased in *Atm*^−/–^ (in which resection usually occurs) and *Mre11*^−/–^ (in which it does not). Otherwise, if the lack of resection enhances end joining, we would observe more frequent deletions in *Mre11*^−/–^. Conversely, if unresected DSBs are not preferred substrates for end ligation, we would observe more events in *Atm*^−/–^. However, it is also possible that unresected and resected ends are joined with different efficiencies, and that the deletion frequencies we observe are influenced by distinct defects in *Mre11*^−/–^ and *Atm*^−/–^, which, together with increased double cuts, promote end joining. For example, MMEJ at resected ends may generally be more active, but the more severe recombination defect in *Mre11*^−/–^ compared with *Atm*^−/–^ may compensate to result in similar numbers of deletions at unresected ends. Nonetheless, although the preferred substrate for end ligation is unclear, we can conclude that abundant double cuts elicit mutagenic end joining.

Whether one end joining pathway predominates or both pathways participate in mutagenic repair remains to be determined. Consistent with some of our results in *Atm*^−/–^ mice^4^ (**Fig. 2D**), MMEJ was hypothesized to mediate deletions found in human population data at single meiotic DSBs, similarly based on the observation of microhomologies at junctions^31^. On the other hand, while NHEJ has been proposed in radiation studies in mice to be active only late in prophase I^60^, after the point of meiotic arrest in both *Mre11*^−/–^ and *Atm*^−/–18,26^, our data argue that NHEJ is also active earlier in meiotic prophase.

At wild-type levels of DSBs, it remains unclear whether MRE11-mediated resection initiation suppresses end joining. *Mre11*^*H129N/*–^ mice were uninformative because they are unexpectedly proficient in resection initiation^18^. These mice, similar to *Nbs1*^*ΔB/ΔB*^ mice, show only a minor defect in SPO11 removal but exhibit reduced resection lengths, demonstrating the role of the MRN complex in resection extension^18^. In this context, our study shows that under-resected ends are not sufficient to promote end joining, at least when resection is reduced on average by half. It is possible that end joining will not occur as long as 3’ overhangs are long enough to mediate strand invasion or bind RPA, which is the case in the *Mre11*^*H129N*^ and *Nbs1*^*ΔB*^ hypomorphs^18,58^. In mitotic cells, RPA prevents annealing between microhomologies, inhibiting MMEJ^61^. It is possible that in *Atm*^−/–^ spermatocytes, where a substantial fraction of ends are also under-resected, the excess of DSBs may deplete the RPA pool, promoting MMEJ^62^.

Considering the mutagenic potential of unresected DSBs, it is important to note that while end joining at unresected double cuts leads to deletions, joining at unresected single DSBs is predicted to be largely error-free due to the presence of 2-bp 5’ overhangs after SPO11 cleavage. However, two single unresected meiotic DSBs may still be aberrantly repaired, for example, to result in translocations. Notably, in *C. elegans* meiosis, chromosome fusions mediated by NHEJ were reported in MRN mutants^32,34,37^.

### MRE11-independent processing of SPO11-DSBs

Endonucleolytic processing of SPO11-bound DSB ends by MRE11 is an early step in homologous recombination. While this essential role of MRE11 is conserved reviewed in^63^, alternative SPO11 removal mechanisms remain unexplored and may differ across species. In this study, we readily recovered end-joining products in the absence of MRE11, revealing the existence of MRE11-independent SPO11 removal in mice. We found that loss of TDP2 reduces end-joining products, indicating that it can partially substitute for MRE11 in removing SPO11. Given that TDP2 converts TOP2-blocked DSBs into NHEJ substrates in mitotic mammalian cells^50,51^, this finding further supports NHEJ activity in *Mre11*^−*/*–^ spermatocytes, with the 2-bp 5’ overhangs likely removed or filled in, although we cannot exclude some minimal end resection by other factors. Interestingly, human TDP2 is unable to remove Spo11 when expressed during meiosis in yeast, which do not have a TDP2 ortholog^52^. The authors suggested that Spo11 is inaccessible to TDP2 in yeast because it is blocked by higher-order protein structures. It is possible that, in mice, TDP2 access to the phosphotyrosyl bonds between SPO11 and DNA, similar to TOP2-DSBs, is facilitated by an accessory protein lacking a yeast ortholog and/or by SPO11 proteolysis^64^.

It remains mysterious how the remaining deletions form in the *Mre11 Tdp2* double mutant. The 5’ flap endonucleases FEN1 and XPG have been implicated in the resolution of TOP2-SSB (single-strand break) complexes in vitro^65^, and FEN1 has also been shown to act on covalent TOP2-DNA complexes in vivo^66^. However, FEN1 and XPG do not act on TOP2-DSBs in vitro and have not been tested *in vivo*^65^, making their role in SPO11-DSB removal unclear. Importantly, MRE11-independent DSB processing involving an unknown mechanism(s) exists in other studied species: In *C. elegans*, deletion of Ku restores resection and recombination in mutants with compromised SPO11 removal by the MRN complex^32,34,37^; in *Tetrahymena thermophila*, DMC1 foci are reduced but still observed in the absence of Mre11 or its cofactor Sae2^39^; and in flies, error-free NHEJ is proposed to occur at DSB ends that underwent resection but remain blocked by Spo11-bound oligos^67^. The alternative SPO11 removal mechanisms and their contributions to normal and mutagenic DSB repair remain to be further elucidated, but the results presented here clearly implicate TDP2 plus at least one other pathway.

Unlike deletions, whose formation in the absence of MRE11partially depends on TDP2, ectopic insertions of double-cut fragments appear to require TDP2. In yeast, double-cut fragments smaller than ∼70 bp are resistant to degradation by Mre11 and therefore have a long lifespan^49^. It may thus be the case that small double-cut fragments are similarly resistant to the TDP2-independent processing mechanism that we infer to be operating on chromosomal DSB ends in MRE11-deficient mice. The mutagenic potential of double-cut fragments may thus depend on both their stability and SPO11 removal. Notably, it is predicted that insertions of double-cut fragments as well as deletions will be absent or much less frequent in yeast due to a reduced dependence on end joining pathways.

### Double cutting within single hotspots in *Mre11*^**–*/*–**^ **and *Atm***^**–*/*–**^

Our finding of frequent microdeletions in *Mre11*^−*/*–^ is consistent with the role of MRE11 in suppressing double cutting within hotspots through ATM activation^4,18,19,25^. When compared to inter-hotspot double cuts, the end-joining profiles at intra-hotspot double cuts were more similar in *Mre11*^−*/*–^ and *Atm*^−*/*–^, indicating that microdeletions occur at unresected or very minimally resected ends in both mutants. Alternatively, joining in *Atm*^−*/*–^ may occur at resected ends but at or very close to the 3′ ends, potentially facilitated by the close proximity of ends.

Nonetheless, we found that microdeletion breakpoints in *Mre11*^−*/*–^ clustered more tightly within the hotspot center than in *Atm*^−*/*–^, suggesting that the double cuts themselves were also more clustered. Our ectopic insert data support this conclusion, as insert source positions in *Atm*^−*/*–^ are more spread out compared to *Mre11*^−*/*–^. While analyses of microdeletions across multiple hotspots are needed to draw definitive conclusions, it is interesting to speculate that the accumulation of MRE11 at hotspot centers in *Atm*^−*/*–18,22^ might discourage SPO11 from catalyzing additional DSBs in close proximity. A non-exclusive possibility is that an MRE11-dependent but ATM-independent chromatin remodeling activity opens the chromatin in cis after DSB formation, allowing additional DSBs to form outside the normal hotspot boundaries in *Atm*^−*/*–^ but not in *Mre11*^−*/*–^.

Notably, our study uncovered potential double cuts of ∼21 bp, which are smaller than the shortest fragments detected in yeast and previously proposed for mice, which were ∼30 bp^4,25,49^. The ∼21-bp microdeletions are evident in *Mre11*^−*/*–^, and their breakpoint distributions within the hotspot support our interpretation that they likely reflect initial SPO11 double-cut positions (**Supplemental Fig. 6D**). Supporting this view, recombinant mouse SPO11 complexes in vitro (unlike the yeast proteins) can simultaneously occupy both ends of a DNA duplex that is just two helical turns long^25,53^. Further, a small fraction of SPO11 oligos <30 nt long, presumably from double cuts, have been observed in *Mre11*^−*/*–^ mice^18^. The frequent occurrence of ∼21-bp microdeletions might indicate that these very short double cuts are joined more efficiently than more widely spaced double cuts.

We also captured even smaller microdeletions, <18 bp, in both *Mre11*^−*/*–^ and *Atm*^−*/*–^. Double cuts of that size are not expected to form, as they are incompatible with the 10-bp periodicity, instead involving ∼1.5 helical turns, and it is unlikely that two adjacent SPO11 complexes could have their active sites that close^25,49,53,56^. Considering these constraints, we envision that <18-bp microdeletions may arise by other mechanisms, e.g., imprecise end joining at single DSBs involving minimal processing and/or partial filling of larger double-cut gaps, particularly in *Atm*^−*/*–^. Alternatively, these events may arise between single DSBs or double cuts formed on both sister chromatids.

## Materials and Methods

### Mice

Mouse experiments were conducted according to US Office of Laboratory Animal Welfare regulatory standards and were approved by the Memorial Sloan Kettering Cancer Center (MSK) Institutional Animal Care and Use Committee (IACUC). Animals were fed standard rodent chow with continuous access to food and water. Previously published *Mre11*^*flox*^ and *Mre11*^*H129N*^ alleles^41^ were crossed with *Ngn3-Cre*^42^ to generate *Mre11*^*flox/flox*^ *Ngn3-Cre+* and *Mre11*^*H129N/flox*^ *Ngn3-Cre+* mice, in which the wild-type allele is conditionally deleted by Cre-mediated recombination in male germ cells^18^. For more efficient *Mre11* knockout in spermatocytes, experiments were performed in *Mre11*^*flox/*–^ *Ngn3-Cre+* animals. *Atm*^68^, *Nbs1*^*ΔB* 59^ and *Tdp2*^51^ mice were previously described and maintained on a predominantly C57BL/6 background. *Atm* mice were purchased from The Jackson Laboratory (B6.129S6-*Atm*^*tm1Awb*^/J; stock #008536) and *Tdp2* mice were a gift from F. Cortés-Ledesma. *Mre11 Tdp2* and *Atm Tdp2* animals were obtained by crossing *Tdp2* mice with *Mre11*^*flox*^ *Ngn3-Cre* or *Atm* mice, respectively.

### Isolation of testis and sperm DNA

Testis DNA was isolated from single-cell suspensions according to the previously published protocol^4^. Briefly, testes were dissected from adult 2–5-month-old mice, except for mice with conditionally deleted *Mre11*, which were 5–7 weeks old to ensure more penetrant Cre-mediated excision^18^. After decapsulation, seminiferous tubules were incubated in 10 ml Gey’s Balanced Salt Solution (GBSS) (Sigma) with 0.5 mg/ml collagenase (Sigma) and 2 μg/ml DNase I (Roche) at 33°C for 15–20 min with shaking (500 rpm). After two washes with 10 ml GBSS, the tubules were digested with 0.5 mg/ml trypsin (Sigma) in 10 ml GBSS containing 2 μg/ml DNase I for 25–40 min at 33°C with shaking (500 rpm). Next, 1 ml fetal calf serum was added, and a cell suspension was obtained by pipetting up and down for 2 min with a plastic transfer pipet, followed by filtration through a 70-µm cell strainer (BD Falcon). The cell suspension was centrifuged for 3 min at 1550 × g, and the cell pellet was resuspended by tapping. Subsequently, 600 μl lysis buffer (200 mM NaCl, 100 mM Tris-Cl pH 7.5, 5 mM EDTA, 0.5% SDS, 0.5 mg/ml Proteinase K (Roche)) was added. The lysate was incubated at 55°C overnight. Phenol/ chloroform/isoamyl alcohol extraction was performed twice, followed by precipitation of the genomic DNA with ice-cold 100% ethanol and two washes with 70% ethanol. DNA was dissolved in 200–400 µl of 5 mM Tris (pH 7.5) overnight at 4°C. DNA was then diluted to ∼200 ng/µl and incubated at 37°C for 2–3 h. After the DNA concentration was measured three times, samples were stored at −20°C.

For experiments with sperm DNA, cauda epididymides were collected from adult mice (2–5 months), and sperm DNA was isolated as previously described^69^.

### Detection of deletions at hotspot pairs by nested PCR

Deletions were detected by nested PCR as previously described (**Fig. 1A**)^4^. Primers flanking the newly assayed hotspot pairs within the Chr13 hotspot cluster (**Supplemental Fig. 1A**) and the published primers for the Chr1 and Chr19 hotspot pairs are listed in **Supplemental Table S1**. Primary and secondary PCR reactions, including S1 digest, were performed in 96-well plates. Each primary 50 µl PCR mix contained: 50 ng input genomic DNA (∼16,000 DNA molecules), 1x buffer (10x: 450 mM Tris-Cl pH 8.8, 110 mM (NH_4_)_2_SO_4_, 45 mM MgCl_2_, 67 mM 2-mercaptoethanol, 44 μM EDTA, 10 mM each dNTP (Roche), 1.13 mg/ml ultra-pure (non-acetylated) BSA (Invitrogen), 12.5 mM Tris base), 0.2 μM primers, 0.03 U/μl Taq DNA polymerase (Thermo Fisher Scientific) and 0.006 U/μl Pfu Turbo DNA polymerase (Agilent). The primary PCR product (0.6 µl) was added to 4.4 µl of the S1 nuclease reaction (0.5× S1 buffer, 0.2 U/μl S1 nuclease (Thermo Fisher Scientific), 5 ng/μl sonicated salmon sperm DNA), incubated at room temperature for 30 min, and then diluted with 45 μl of dilution buffer (10 mM Tris-Cl pH 7.5 and 5 ng/μl sonicated salmon sperm DNA). The diluted S1 reaction (4 μl) was used to seed a 20 µl secondary PCR reaction, which had the same composition as the primary PCR.

Conditions for the primary and secondary PCR were the same: denaturation (2 min at 96°C), followed by 30 cycles of amplification (denaturation for 20 s at 96°C, annealing for 30 s at 61°C for the Chr13 hotspot pairs and 59°C for the Chr1 and Chr19 hotspot pairs, extension at 65°C for 4 min for the Chr13 hotspot pairs and 2 min 30 s for the Chr1 and Chr19 hotspot pairs). For the Chr13 hotspot pairs, the 4 min extension time allowed amplification of only deletion products, not parental products (**Supplemental Table S1**). However, for the 5 kb hotspot pair, a 4.2 kb PCR artifact was detected in addition to smaller deletion products (**Supplemental Fig. 1B**). For the Chr1 and Chr19 hotspot pairs, 2 min 30 s extension, which is shorter than the optimal <60 s/kb, helped inhibit amplification of the 3.4-kb parental products^4^.

Deletion products were detected by electrophoresis on a 1% agarose gel (**Fig. 1B**), excised and purified (Invitrogen PureLink™ Quick Gel Extraction Kit), and Sanger sequenced. Sequenced PCR products were analyzed with ApE software (https://jorgensen.biology.utah.edu/wayned/ape/) to map deletion breakpoints and identify microhomologies. Downstream analyses, including deletion breakpoint profiling, were performed using R version 4.2.2 to 4.5.2 (http://www.r-project.org). Microhomology expected by chance was calculated as the probability of microhomology at a junction of a given length (x) in a random DNA sequence, using the equation P(x) = (x + 1)(1/4)^x^(3/4)^2 70^. Graphing and statistical analyses were performed in R and GraphPad Prism version 10.

### Detection of microdeletions at a single hotspot by amplicon deep sequencing

Generation of amplicons and deep sequencing were performed as previously described^4^. Briefly, four overlapping ∼580-bp amplicons of the Chr1 single hotspot (**Supplemental Fig. 5A**) were amplified using published primers^4^. PCRs were carried out in 96-well plates in 50 µl reactions, each containing 50 ng input genomic DNA, 1.5 U Taq DNA polymerase (Thermo Fisher Scientific), and 0.3 U Pfu Turbo DNA polymerase (Agilent) in the buffer described above for nested PCR. For each amplicon, PCR conditions were the same: denaturation (2 min at 96°C), followed by 22 cycles of amplification (20 s at 96°C, 30 s at 59°C, and 35 s at 65°C). Testis DNA was isolated from three *Mre11*^−*/*–^ mice and amplified independently. A total of 576 PCRs were performed for each amplicon, with input DNA equivalent to 9.6 million haploid genomes. The high number of PCRs combined with low amplification cycles, which minimize PCR bias, generates amplicons with large numbers of independently amplified products for sequencing. Next, for each amplicon, PCR products were pooled, and a 1200-μl aliquot was purified and concentrated on spin columns (Invitrogen PureLink™ PCR Purification Kit), as described in detail previously^4^. Approximately 60 µl of DNA at a concentration of 1–2 ng/µl was submitted to the Integrated Genomics Operation at MSK for library preparation and deep sequencing. Indexed libraries were prepared using the KAPA HyperPrep Kit (Roche) and sequenced on the Illumina MiSeq platform, generating paired-end reads of 300 bp. The four amplicon libraries were sequenced in a single run with a 5% PhiX DNA spike-in, estimating 3.6–4.8 million reads per amplicon. Given the 9.6 million input DNA molecules, most captured deletions should be represented by a single read.

The obtained raw sequencing data were analyzed using a new bioinformatic pipeline, svAmplicon, an extension of the svx-mdi-tools suite repository^71^, since the previously used CRISPResso2 was retired^4^. Paired-read FASTQ sequencing files were quality filtered and merged when sufficiently overlapping into single reads by fastp version 0.23.2. Prepared reads were aligned to the mm10 reference genome using bwa mem version 0.7.17 (https://arxiv.org/abs/1303.3997) to identify the best match for each read while allowing for off-target alignment. The outer endpoints of reads were used to perform *ab initio* identification of apparent amplicons present in the pooled data, which consistently met expectations of the PCR reactions performed. Reads were grouped into unique sequences per amplicon, keeping the best base quality observed over all equivalent reads, and realigned to the genome as individual best. Structural variant (SV) breakpoint nodes were first identified from discontinuous, i.e. supplementary alignments, and then by extracting small inline node pairs from the CIGAR string of individual alignments representing deletions 10 bp or larger. An important class of artifactual false junctions can arise when a low-quality segment of a read fails to align in an otherwise continuous amplicon, leading to an insertion and deletion of roughly equal length. Such non-SV indels were identified and discarded, resulting in called SVs with a net deletion of ≥10 bp. Read sequences were parsed to infer the microhomologous or inserted bases at each junction^72^. Amplicon reads were grouped into unique alignment paths to allow potential PCR duplicates to be identified. Final junction validation was performed in the interactive svAmplicon R Shiny app. First, filters required that the best read representing a molecule path had an average Phred-scaled base quality of 25 and a mapping quality (MAPQ) of 55 for each alignment. Deletion SVs as defined above were individually examined, allowing correlation of local base quality, indel lengths, and microhomology spans to ensure that only true novel junctions were tabulated in **Dataset S1**. We also reanalyzed our published *Atm*^−*/*–^ data^4^. Although we observed a higher total number of unique (independent) microdeletions in *Mre11*^−*/*–^ (**Supplemental Table S8**), the better sequencing quality of *Mre11*^−*/*–^ amplicons and the different yields between batches hinder an accurate quantitative comparison.

Downstream analyses of unique microdeletions and plotting were performed using R (versions 4.2.2–4.5.2). For consistency with published results, microdeletion breakpoint positions and sizes were adjusted (**Dataset S1**).

### Data, Materials, and Software Availability

Raw amplicon sequencing *Mre11*^−*/*–^ data will be deposited in the Gene Expression Omnibus (GEO) repository. Processed amplicon sequencing data from *Mre11*^−*/*–^ and from reanalyzed *Atm*^−*/*–^ (GEO accession number GSE182210)^4^ are provided in **Dataset S1**. The svAmplicon pipeline and app code are available as part of the svx-mdi-tools repository (GitHub: https://github.com/wilsontelab/svx-mdi-tools; documentation: https://wilsontelab.github.io/svx-mdi-tools), linked to Zenodo DOI (https://doi.org/10.5281/zenodo.7871676). The job files used to execute the pipeline for *Mre11*^−*/*–^ and *Atm*^−*/*–^ samples and associated log reports are available as **Dataset S2**, which, together with the installed code, provides complete details for repeating the job execution. We used the *Atm*^−*/*–^ dataset “*Atm* null 1” from SPO11-oligo data (GEO accession number GSE84689)^8^. Original data including gel pictures, Sanger sequencing, and any additional information required to reanalyze the data reported in this study are available from the lead contact upon request.

## Acknowledgments

We thank Felipe Cortés-Ledesma (CNIO) for providing *Tdp2* mutant mice; members of the Lukaszewicz, Jasin, and Keeney labs for comments and discussions; and MSK core facilities, in particular the Integrated Genomics Operation, which are supported by NIH (P30 CA008748). This work was supported by NIH grants R35 GM157077 (A.L.), R35 GM118092 (S.K.), and R01 HD112624 (M.J.).

This article is subject to HHMI’s Open Access to Publications policy. HHMI lab heads have previously granted a nonexclusive CC BY 4.0 license to the public and a sublicensable license to HHMI in their research articles. Pursuant to those licenses, the author-accepted manuscript of this article can be made freely available under a CC BY 4.0 license immediately upon publication.

## Author contributions

A.L., S. Keeney, and M.J. conceived the project. S. Kim provide unpublished data and critical reagents. A.L. performed the experiments. A.L. and T.W. analyzed the data. A.L. and M.J. wrote the manuscript with input from S. Keeney and T.W.

