## Supplement for "MRE11 suppresses germline mutagenesis at meiotic double-strand breaks in mice"

**This pdf contains:**

Supplemental Figures 1–7
Supplemental Tables 1–8

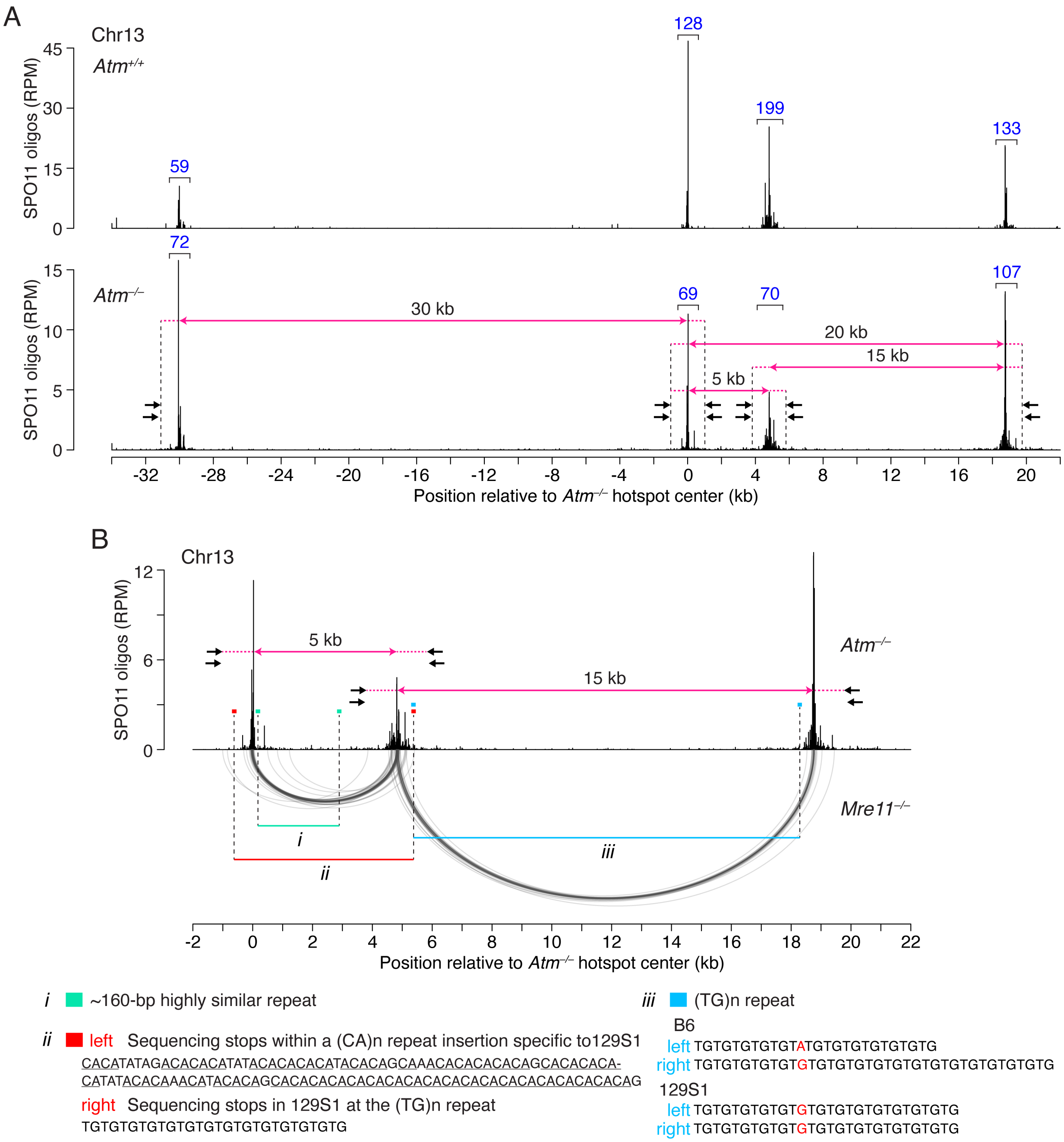


**Supplemental Figure S1. Deletions at the Chr13 hotspot cluster.**
**A.** SPO11-oligo maps for the hotspot cluster on Chr13 in wild-type and *Atm*$^{-/-}$[8]. Numbers in blue denote hotspot strength in reads per million (RPM) and do not take into account the overall 11.3-fold higher number of SPO11-oligos in *Atm*$^{-/-}$[19]. The increase in hotspot strength is not uniform across hotspots, but rather weaker hotspots increase proportionately more than stronger hotspots[19]. The 0-kb position is arbitrarily assigned to the second hotspot.

The PCR design is shown on the *Atm*$^{-/-}$ map. Nested PCR primers (arrows) used to detect deletions at the hotspot pairs are shown together with the distances between hotspot centers (pink horizontal lines). Regions (dashed pink horizontal lines) between hotspot centers and the corresponding flanking secondary PCR primers (dashed black vertical lines) are indicated.

**B.** Bona fide deletion events (shown as arcs below the SPO11-oligo maps) and presumptive PCR artifacts at the 5-kb and 15-kb hotspot pairs. Three deletion types (*i, ii,* and *iii)* considered to be PCR artifacts arise at the indicated repetitive sequences and were excluded from subsequent analyses (**Supplemental Tables S2–S5**). Explanations for why they are presumably artifacts are below. The deletion maps shown are from *Mre11*$^{-/-}$ mice; the artifacts can be seen in *Atm*$^{-/-}$ as well.

*i.* 5-kb hotspot pair: The parental PCR product for the 5-kb hotspot pair (6.9 kb) is not expected to be amplified, but a smaller PCR product (4.2 kb) was detected from each PCR. This PCR product contains a deletion between two ~160-bp highly similar repeats and is observed in every genotype tested, so it is likely a PCR artifact. This artifact does not impede detection of deletions, as the right repeat is located between the two hotspots. It is unlikely that we are missing smaller deletions with both breakpoints mapping between the two repeats because there are no observed right breakpoints mapping close to the right repeat.

*ii.* 5-kb hotspot pair: This deletion product arises only when the Chr13 hotspot cluster is from the 129S1 strain background. While both the 129S1 and B6 Chr13 hotspot clusters have a $(TG)_n$ repeat located near the right hotspot, the 129S1 sequence additionally contains an insertion of a $(CA)_n$ repetitive sequence (underlined) near the left hotspot. Deletions between the $(CA)_n$ and $(TG)_n$ repeats could not be fully sequenced because sequencing stops within the two repeats. Given the repetitive sequences near the breakpoints and the detection of these events at similar frequencies when tested side by side in *Mre11*$^{-/-}$ and *Mre11*$^{+/-}$ (**Supplemental Table S3**), we consider them to be PCR artifacts.

*iii*. 15-kb hotspot pair: This deletion product involves two $(TG)_n$ repeats which are identical when the Chr13 hotspot cluster is from the 129S1 strain background but differ when the cluster is from the B6 strain background. Deletions are rare on the B6 background, but because they can be found even in *Spo11*$^{-/-}$ they are excluded from subsequent analyses. In 129S1, these deletions at the identical repeats are more frequent, such that they can impact the frequencies of bona fide deletions. Thus, we only analyzed deletions at the 15-kb hotspot when the hotspot clustered was homozygous for B6.

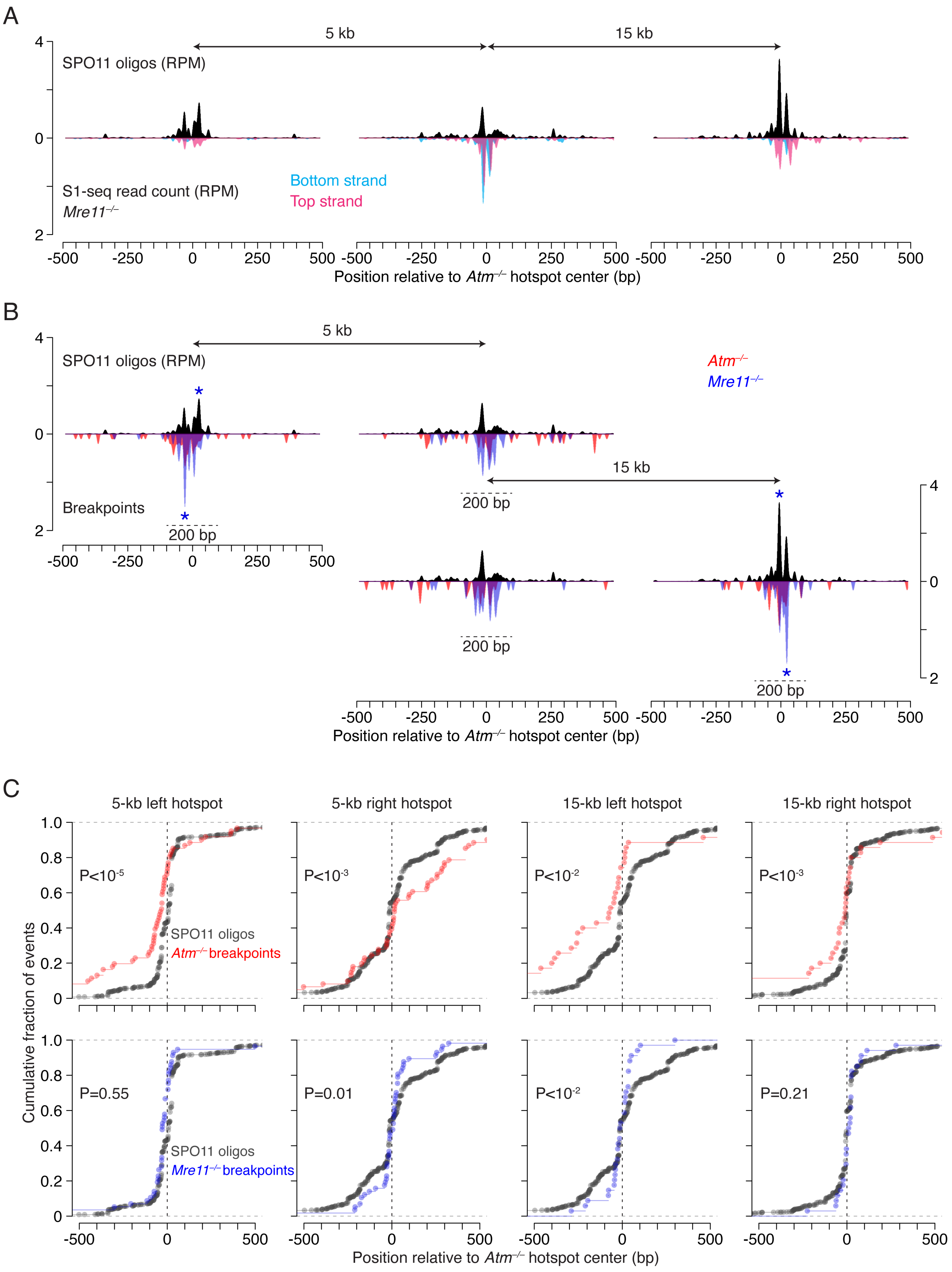

A
5 kb
15 kb
SPO11 oligos (RPM)
S1-seq read count (RPM)
Mre11−/−
Bottom strand
Top strand
Position relative to Atm−/− hotspot center (bp)
B
5 kb
15 kb
SPO11 oligos (RPM)
Atm−/−
Mre11−/−
Breakpoints
200 bp
Position relative to Atm−/− hotspot center (bp)
C
5-kb left hotspot
5-kb right hotspot
15-kb left hotspot
15-kb right hotspot
P<10-5
P<10-3
P<10-2
P<10-3
SPO11 oligos
Atm−/− breakpoints
P=0.55
P=0.01
P<10-2
P=0.21
SPO11 oligos
Mre11−/− breakpoints
Cumulative fraction of events
Position relative to Atm−/− hotspot center (bp)

**Supplemental Figure S2. Comparison of deletion breakpoint distributions in *Atm*$^{-/-}$ and *Mre11*$^{-/-}$.**

**A.** SPO11-oligo and *Mre11*$^{-/-}$ strand-specific S1-seq profiles[18] at the 5-kb and 15-kb hotspot pairs. For S1-seq, signals from two biological replicate libraries were averaged and plotted around SPO11-oligo hotspot centers. Plots are smoothed with a 21-bp Hann filter.

**B.** Smoothed breakpoint and SPO11-oligo profiles (21-bp Hann filter) for the 5-kb and 15-kb hotspot pairs. Breakpoint profiles are normalized to the number of deletions in *Atm*$^{-/-}$ at the 5-kb hotspot pair. Dashed line indicates a 200-bp window around the hotspot center. Blue asterisks mark offset major *Mre11*$^{-/-}$ breakpoint and SPO11-oligo peaks, consistent with double cuts at the major and secondary SPO11-oligo peaks.

**C.** Cumulative fractions of SPO11 oligos and *Atm*$^{-/-}$ or *Mre11*$^{-/-}$ breakpoints around hotspot centers. For SPO11-oligo distributions, only reads mapping within ±1500 bp from hotspot centers were included. P values are from Levene's test.

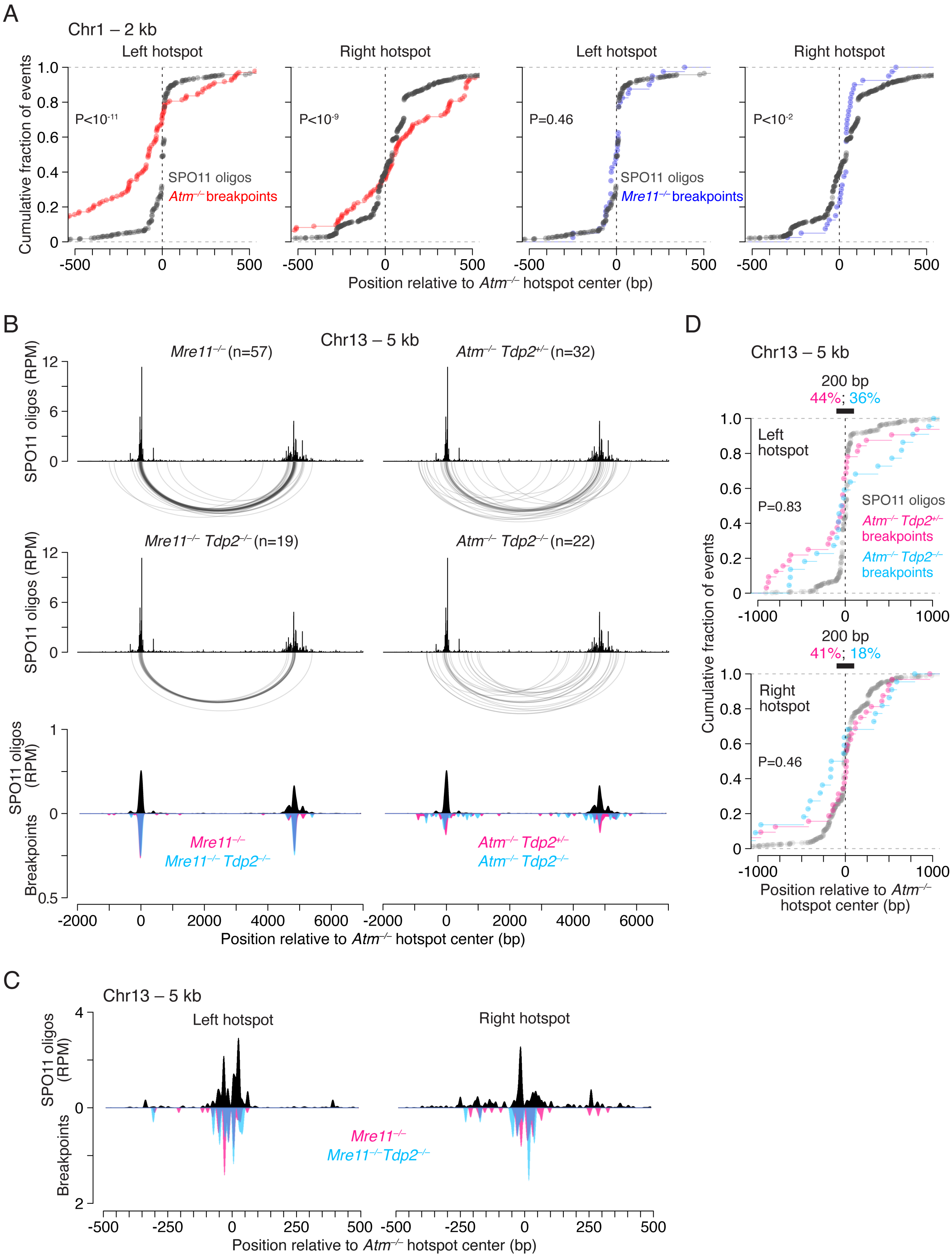

A
Chr1 – 2 kb
Left hotspot
Right hotspot
Left hotspot
Right hotspot
Cumulative fraction of events
P<10-11
P<10-9
P=0.46
P<10-2
SPO11 oligos
Atm-/- breakpoints
SPO11 oligos
Mre11-/- breakpoints
Position relative to Atm-/- hotspot center (bp)
B
Chr13 – 5 kb
Mre11-/- (n=57)
Atm-/- Tdp2+/- (n=32)
Mre11-/- Tdp2-/- (n=19)
Atm-/- Tdp2-/- (n=22)
SPO11 oligos (RPM)
SPO11 oligos (RPM)
Breakpoints
Mre11-/-
Mre11-/- Tdp2-/-
Atm-/- Tdp2+/-
Atm-/- Tdp2-/-
Position relative to Atm-/- hotspot center (bp)
C
Chr13 – 5 kb
Left hotspot
Right hotspot
SPO11 oligos (RPM)
Breakpoints
Mre11-/-
Mre11-/-Tdp2-/-
Position relative to Atm-/- hotspot center (bp)
D
Chr13 – 5 kb
200 bp
44%; 36%
Left hotspot
P=0.83
SPO11 oligos
Atm-/- Tdp2+/- breakpoints
Atm-/- Tdp2-/- breakpoints
200 bp
41%; 18%
Right hotspot
P=0.46
Cumulative fraction of events
Position relative to Atm-/- hotspot center (bp)

**Supplemental Figure S3. Impact of MRE11 and TDP2 on end joining.**
**A.** Cumulative fractions of SPO11 oligos and *Atm*$^{-/-}$ or *Mre11*$^{-/-}$ breakpoints around the hotspot centers of the 2-kb hotspot pair on Chr1. *Atm*$^{-/-}$ data are from the previous study[4]. For SPO11-oligo plots, reads mapping within ±1500 bp from hotspot centers were included. P values are from Levene's test.
**B.** Comparisons of deletion breakpoint distributions with and without TDP2 in MRE11- or ATM-deficient backgrounds. Profiles at the bottom are smoothed with a 151-bp Hann filter and normalized to the number of events in *Atm*$^{-/-}$ *Tdp2*$^{+/-}$.
**C.** Smoothed breakpoint and SPO11-oligo plots (21-bp Hann filter) for the 5-kb hotspot pair on Chr13. *Mre11*$^{-/-}$ and *Mre11*$^{-/-}$ *Tdp2*$^{-/-}$ breakpoint profiles overlap near the hotspot centers. Breakpoints are normalized to the number of events in *Mre11*$^{-/-}$.
**D**. Cumulative distributions of end-joining breakpoints in *Atm*$^{-/-}$ mutants with (*Tdp2*$^{+/-}$) and without (*Tdp2*$^{-/-}$) compared to SPO11 oligos, around the left and right hotspot of the 5-kb hotspot pair on Chr13. For SPO11 oligos, reads mapping within ±1500 bp from hotspot centers were included. Percentages above the plots indicate the fraction of breakpoints within a 200-bp window (black bar) around the hotspot centers. P values are from Levene's test comparing breakpoint distributions between the two genotypes.

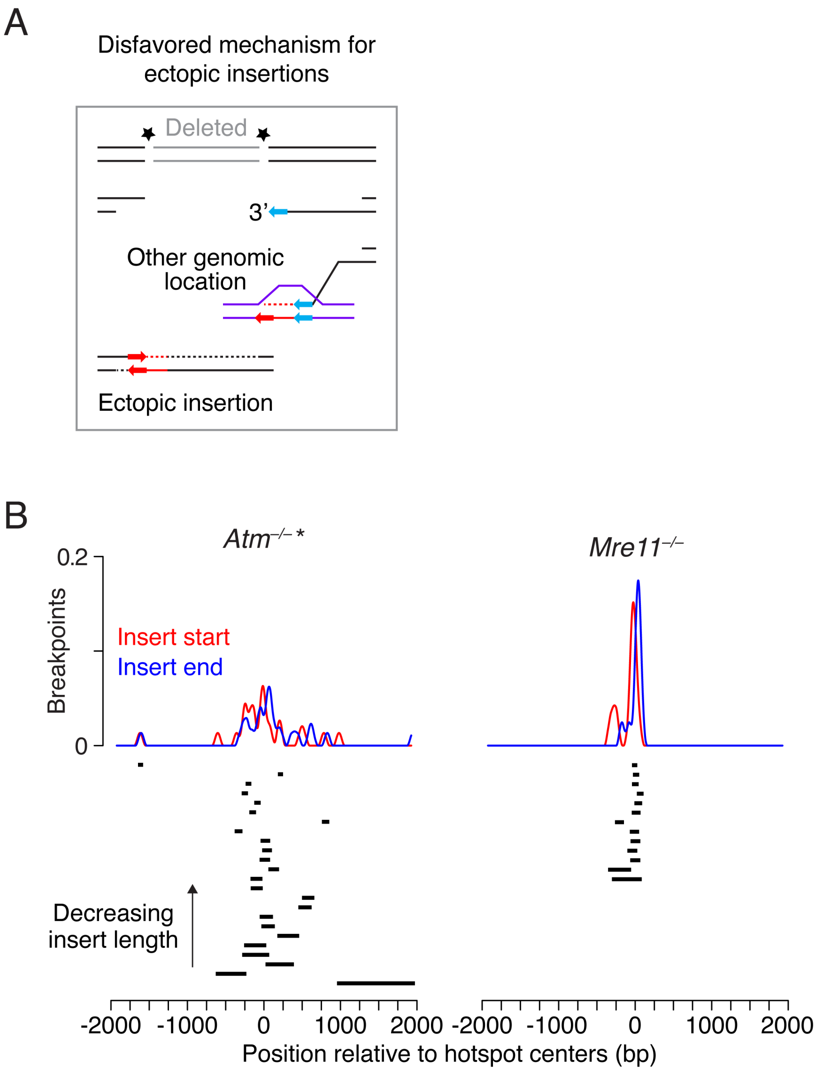


**Supplemental Figure S4. Ectopic insertions in the absence of MRE11.**
**A.** Alternative scenario from **Fig. 4A** for generation of deletions with ectopic insertions, in which a resected end invades and copies a non-allelic template, then anneals back to the other resected DSB end. This mechanism is promoted by microhomologies (arrows). In contrast to ectopic insertion of double-cut fragments (**Fig. 4A**), this synthesis-based scenario should not preferentially involve other hotspots.
**B.** Inserts from other hotspots mapped relative to their respective hotspot centers and sorted by length. Smoothed insert breakpoint profiles (151-bp Hann filter) are normalized to the number of events in *Atm*$^{-/-}$. *Atm*$^{-/-}$ * indicates combined *Atm*$^{-/-}$ and *Atm*$^{-/-}$ *Tdp2*$^{+/-}$ inserts, including previous *Atm*$^{-/-}$ data[4]. Three inserts in *Atm*$^{-/-}$ mapping to multiple hotspots on non-PAR ChrY were excluded here and in **Fig. 4D**[4].

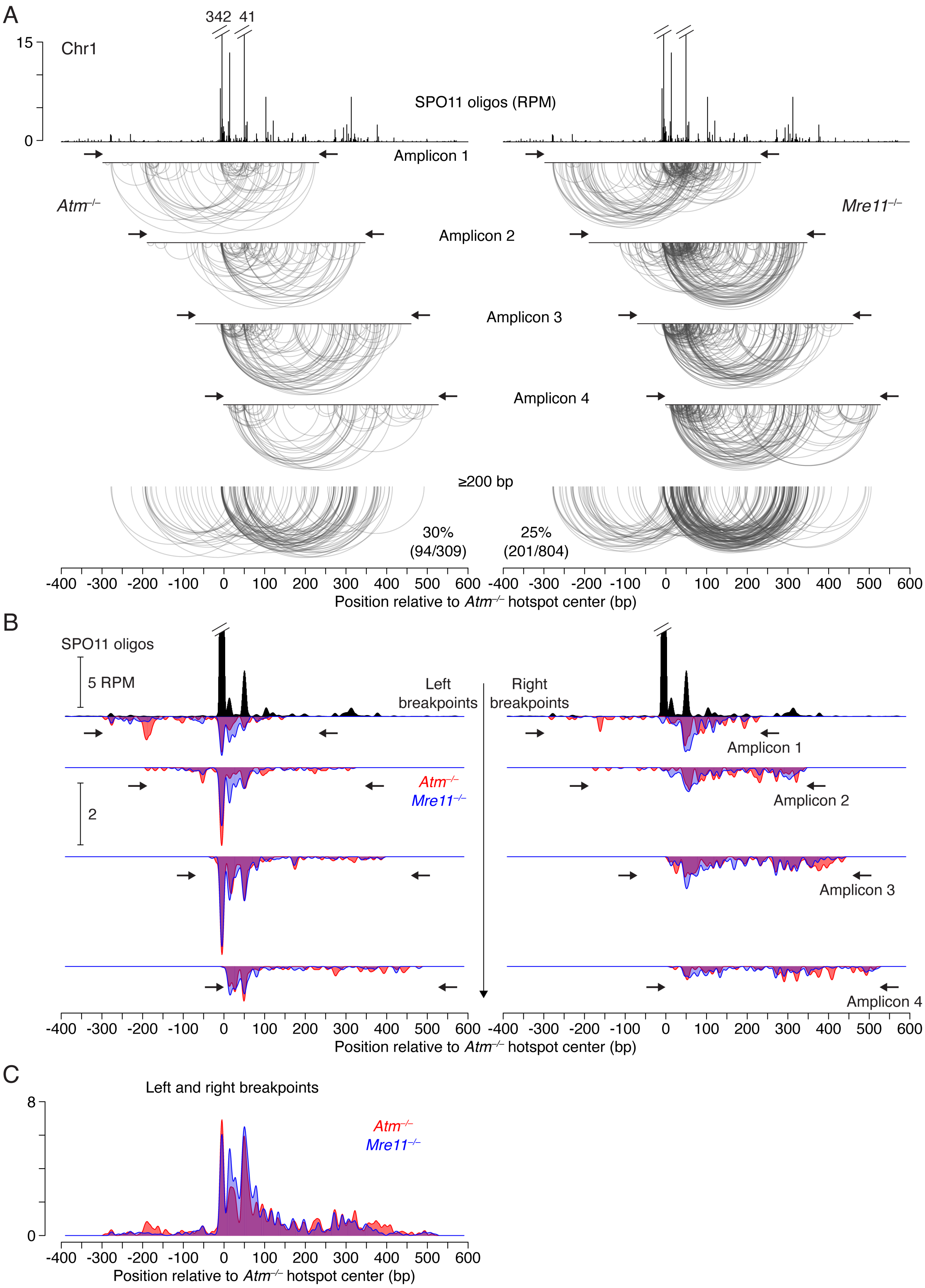
A
342
41
15
0
Chr1
SPO11 oligos (RPM)
Amplicon 1
Atm−/−
Mre11−/−
Amplicon 2
Amplicon 3
Amplicon 4
≥200 bp
30%
(94/309)
25%
(201/804)
-400 -300 -200 -100 0 100 200 300 400 500 600
Position relative to Atm−/− hotspot center (bp)
B
SPO11 oligos
5 RPM
Left
breakpoints
Right
breakpoints
Amplicon 1
Atm−/−
Mre11−/−
2
Amplicon 2
Amplicon 3
Amplicon 4
-400 -300 -200 -100 0 100 200 300 400 500 600
Position relative to Atm−/− hotspot center (bp)
C
Left and right breakpoints
8
0
Atm−/−
Mre11−/−
-400 -300 -200 -100 0 100 200 300 400 500 600
Position relative to Atm−/− hotspot center (bp)

**Supplemental Figure S5. Analyses of microdeletions at a single hotspot in *Mre11*$^{-/-}$ and *Atm*$^{-/-}$.**

**A.** Distributions of independent microdeletions (11 bp to 361 bp) found in each amplicon for *Mre11*$^{-/-}$ and *Atm*$^{-/-}$, relative to the SPO11-oligo distribution. Arrows indicate primers used to amplify ~580 bp of the hotspot region in the four amplicons. Below are microdeletions ≥200 bp, filtered out from the four amplicons, with the indicated percentage of all events. The previously published *Atm*$^{-/-}$ amplicons[4] were reanalyzed. For microdeletion frequencies per amplicon, see **Supplemental Table S8**.

**B.** Comparison of smoothed left and right breakpoint maps between *Mre11*$^{-/-}$ and *Atm*$^{-/-}$, relative to SPO11-oligo maps (21-bp Hann filter), for each amplicon (arrows indicate primers). Breakpoints are normalized to *Atm*$^{-/-}$ events.

**C.** End-joining profiles in *Mre11*$^{-/-}$ and *Atm*$^{-/-}$ for combined left and right breakpoints. Profiles are smoothed (21-bp Hann filter) and normalized to *Atm*$^{-/-}$ events.

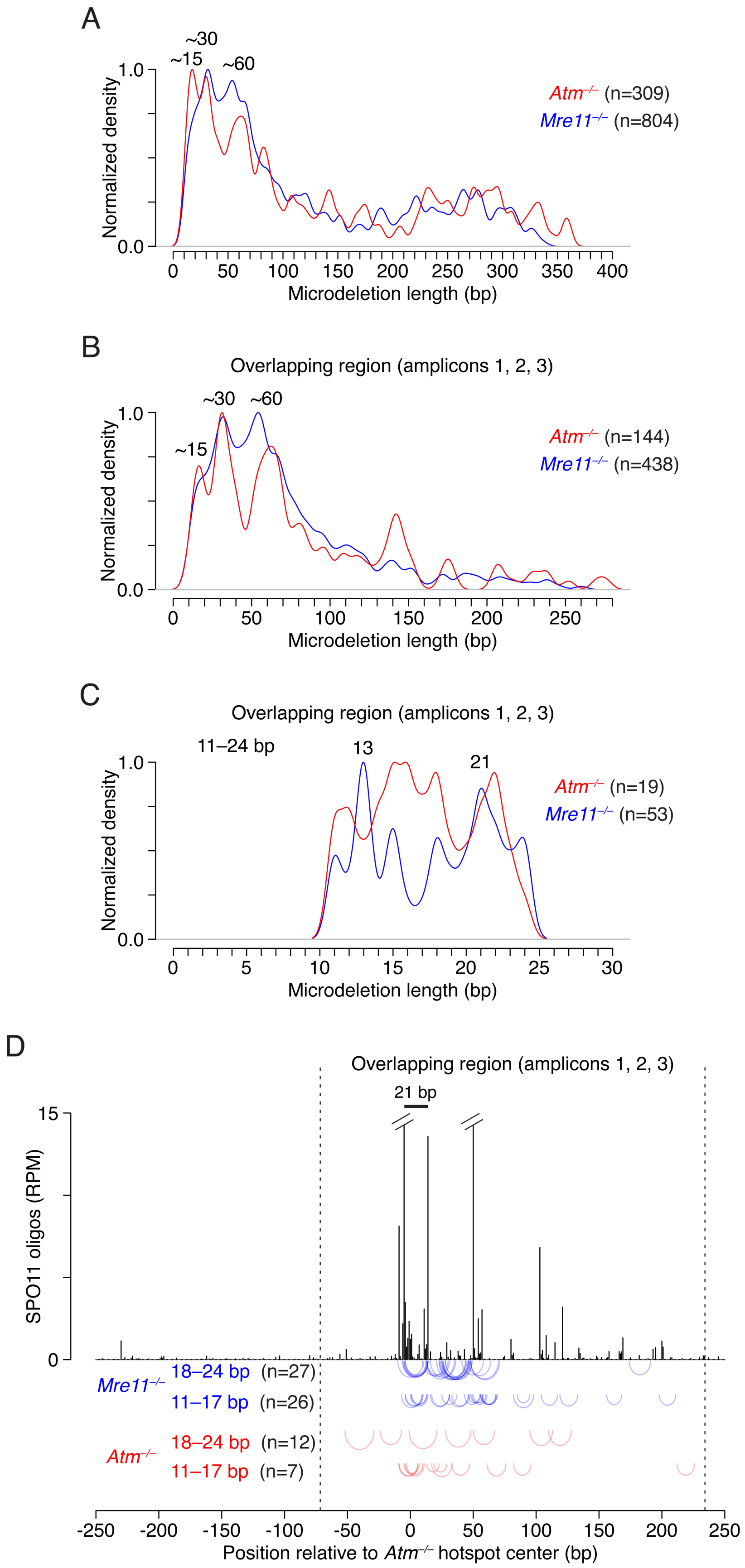
A
~30
~15
~60
1.0
0.0
Normalized density
Atm−/− (n=309)
Mre11−/− (n=804)
0
50
100
150
200
250
300
350
400
Microdeletion length (bp)
B
Overlapping region (amplicons 1, 2, 3)
~30
~60
~15
1.0
0.0
Normalized density
Atm−/− (n=144)
Mre11−/− (n=438)
0
50
100
150
200
250
Microdeletion length (bp)
C
Overlapping region (amplicons 1, 2, 3)
11–24 bp
13
21
1.0
0.0
Normalized density
Atm−/− (n=19)
Mre11−/− (n=53)
0
5
10
15
20
25
30
Microdeletion length (bp)
D
Overlapping region (amplicons 1, 2, 3)
21 bp
15
0
SPO11 oligos (RPM)
Mre11−/−
18–24 bp (n=27)
11–17 bp (n=26)
Atm−/−
18–24 bp (n=12)
11–17 bp (n=7)
-250
-200
-150
-100
-50
0
50
100
150
200
250
Position relative to Atm−/− hotspot center (bp)

**Supplemental Figure S6. Analyses of small microdeletions in *Mre11*$^{-/-}$ and *Atm*$^{-/-}$.**
**A,B.** Length distributions of microdeletions (>10 bp) from all amplicons (**A**) and from the overlapping region of amplicons 1, 2, and 3 (**B**).
**C.** Fine-scale length distributions of microdeletions that are roughly ~15-bp long.
**D.** Distributions of microdeletions (arc diagrams) in the indicated size classes, chosen to match preferred sizes in *Mre11*$^{-/-}$ in panel **C**. The distance between the major and adjacent SPO11-oligo peaks is indicated.

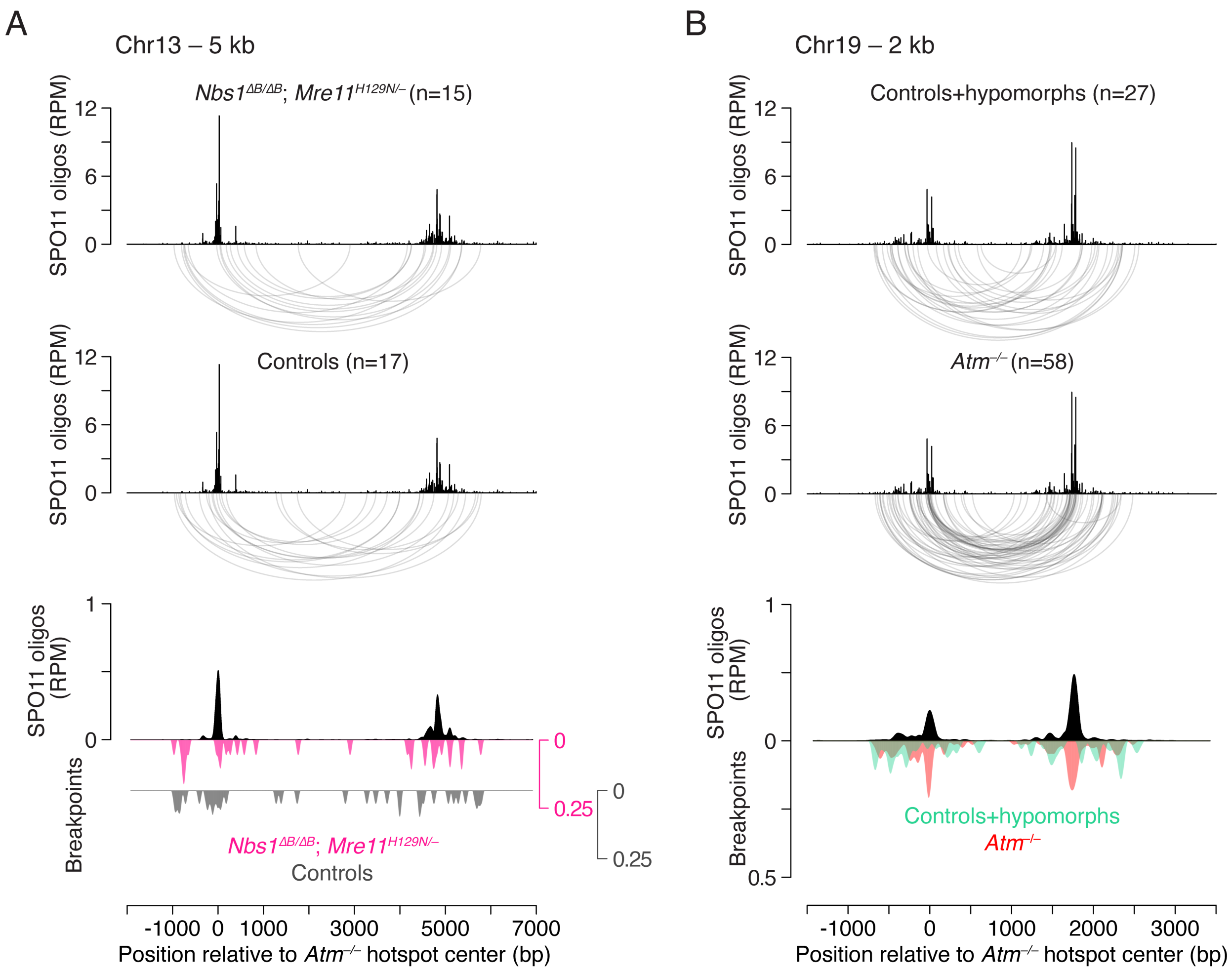


**Supplemental Figure S7. Analyses of deletion breakpoints in *Mre11*$^{H129N/–}$ and *Nbs1*$^{\Delta B/\Delta B}$ hypomorphic mice.**

**A.** Deletion breakpoint distributions in *Mre11*$^{H129N/–}$ and *Nbs1*$^{\Delta B/\Delta B}$ are similar to controls at the 5-kb hotspot pair. Controls are from wild-type and heterozygous mice, including events from other experiments. Breakpoints are normalized to *Atm*$^{–/–}$ deletions. Maps are smoothed with a 151-bp Hann filter.

**B.** Broader deletion breakpoint distributions in *Nbs1*$^{\Delta B/\Delta B}$ and controls at the 2-kb hotspot pair than in *Atm*$^{–/–}$. Events in controls represent deletions shown in **Fig. 6B** and *Atm*$^{+/–}$ and *Atm*$^{+/+}$ deletions found here and in the previous study[4]. *Atm*$^{–/–}$ deletions were previously published[4]. Breakpoint and SPO11-oligo maps are smoothed (151-bp Hann filter), and breakpoints are normalized to *Atm*$^{–/–}$.

### Supplemental Table S1. Primers used to detect deletions at hotspot pairs

| Hotspot pair | Distance between hotspot centers (kb) | Primer[a] | Sequence (5' to 3') | GRCm38/mm10 | Parental 1º and 2º PCR products (kb) | 2º PCR products with deletions at hotspot centers (kb)[b] |
|---|---|---|---|---|---|---|
| Chr13 – 5 kb | 4.8 | Chr13_5kb_FW1 | ACCTCCAAAAAGCTCTGTCTC | chr13:12498546 | 1º PCR – 7.0 | 2.1 |
| | | Chr13_5kb_RV1 | TGTGCTTGGCAGGTTTCCAG | chr13:12505530 | | |
| | | Chr13_5kb_FW2 | GACTCATTTAGCAGAGCATCC | chr13:12498586 | 2º PCR – 6.9 | |
| | | Chr13_5kb_RV2 | CTGTTCCTCATCTAGTGCTTG | chr13:12505474 | | |
| Chr13 – 15 kb | 13.9 | Chr13_15kb_FW1 | TGTCAGGGATTTGGTCTTCGC | chr13:12503357 | 1º PCR – 16.2 | 2.1 |
| | | Chr13_20kb_RV1 | CCTGGTTGAAACAAGTCAGCC | chr13:12519519 | | |
| | | Chr13_15kb_FW2 | TCCCCAGTTTCGTGGAAGTC | chr13:12503401 | 2º PCR – 16.0 | |
| | | Chr13_20kb_RV2 | CTTGTTCGTCTTCTCCCTCC | chr13:12519413 | | |
| Chr13 – 20 kb | 18.8 | Chr13_5kb_FW1 | ACCTCCAAAAAGCTCTGTCTC | chr13:12498546 | 1º PCR – 21.0 | 2.1 |
| | | Chr13_20kb_RV1 | CCTGGTTGAAACAAGTCAGCC | chr13:12519519 | | |
| | | Chr13_5kb_FW2 | GACTCATTTAGCAGAGCATCC | chr13:12498586 | 2º PCR – 20.8 | |
| | | Chr13_20kb_RV2 | CTTGTTCGTCTTCTCCCTCC | chr13:12519413 | | |
| Chr13 – 30 kb | 30.0 | Chr13_30kb_FW1 | CAAGCATTATCTGCAAGCCTC | chr13:12468359 | 1º PCR – 32.4 | 2.2 |
| | | Chr13_30kb_RV1 | TTCTTCTACGATCTTCTCTCAGAC | chr13:12468496 | | |
| | | Chr13_30kb_FW2 | CCCAATTTTACCACCATCGAG | chr13:12500716 | 2º PCR – 32.2 | |
| | | Chr13_30kb_RV2 | AGAGTGGAGTGTGTGTGTTCC | chr13:12500678 | | |
| Chr1 – 2 kb[c] | 1.8 | Chr1_DH_FW1 | GGTCAGGAGGATGGGGAG | chr1:164441295 | 1º PCR – 3.6 | 1.6 |
| | | Chr1_DH_RV1 | CCAAGAAGGTGCTTATCGTCC | chr1:164444852 | | |
| | | Chr1_DH_FW2 | GCCTTGGCTCAAAGTGTCTG | chr1:164441326 | 2º PCR – 3.4 | |
| | | Chr1_DH_RV2 | TCCCTTCTGCCATCCTTCTG | chr1:164444763 | | |
| Chr19 – 2 kb[c] | 1.8 | Chr19_DH_FW2 | TGGAATGTAGCCTCTGGCAG | chr19:24380779 | 1º PCR – 3.6 | 1.6 |
| | | Chr19_DH_RV2 | CCCAAAGTTGTCCTTGCTTCC | chr19:24380968 | | |
| | | new_Chr19_DH_FW2 | GTGCTGTCAGAAGCCGTGG | chr19:24380948 | 2º PCR – 3.4 | |
| | | new_Chr19_DH_RV2 | TGGTTTCCAGAGGAGAGGAC | chr19:24384316 | | |

[a] Primers for 1º and 2º PCR reactions are grouped.
[b] Double cutting at two hotspot centers with precise end joining.
[c] Primers used previously[4].

**Supplemental Table S2. Frequency of deletions in ATM-proficient and -deficient testes**

| Hotspot pair | Mouse and genotype | No. haploid genomes | No. deletions | Frequency ($10^{-6}$) |
|---|---|---|---|---|
| Chr13 – 5 kb | 3777 $Atm^{-/-}$ | 800000 | 2 | 2.50 |
| | 3795 $Atm^{-/-}$ | 800000 | 7 | 8.75 |
| | 3793 $Atm^{+/-}$ | 800000 | 0 | <1.25 |
| | 3835 $Atm^{-/-}$ | 800000 | 8 | 10.00 |
| | 3833 $Atm^{+/-}$ | 800000 | 0 | <1.25 |
| | 3834 $Atm^{+/+}$ | 800000 | 0 | <1.25 |
| | 3849 $Atm^{-/-}$ | 800000 | 11 | 13.75 |
| | 3850 $Atm^{+/-}$ | 800000 | 0 | <1.25 |
| | 3991 $Atm^{-/-}$ | 800000 | 1 | 1.25 |
| | 4286 $Atm^{-/-}$ | 800000 | 12 | 15.00 |
| | 4288 $Atm^{+/+}$ | 1600000 | 1 | 0.63 |
| | Total $Atm^{-/-}$ | 4800000 | 41 | 8.54 |
| | Total $Atm^{+/-}$; $Atm^{+/+}$ | 4800000 | 1 | 0.21 |
| Chr13 – 15 kb | 3777 $Atm^{-/-}$ | 800000 | 2 | 2.50 |
| | 3795 $Atm^{-/-}$ | 800000 | 5 | 6.25 |
| | 3793 $Atm^{+/-}$ | 800000 | 0 | <1.25 |
| | 3835 $Atm^{-/-}$ | 800000 | 3 | 3.75 |
| | 3833 $Atm^{+/-}$ | 800000 | 0 | <1.25 |
| | 3834 $Atm^{+/+}$ | 800000 | 0 | <1.25 |
| | 3849 $Atm^{-/-}$ | 800000 | 8 | 10.00 |
| | 3850 $Atm^{+/-}$ | 800000 | 0 | <1.25 |
| | 3991 $Atm^{-/-}$ | 800000 | 4 | 5.00 |
| | 4286 $Atm^{-/-}$ | 800000 | 13[a] | 16.25 |
| | 4288 $Atm^{+/+}$ | 1600000 | 0[a] | <1.25 |
| | Total $Atm^{-/-}$ | 4800000 | 35 | 7.29 |
| | Total $Atm^{+/-}$; $Atm^{+/+}$ | 4800000 | 0 | <0.21 |
| Chr13 – 20 kb | 3777 $Atm^{-/-}$ | 800000 | 0 | <1.25 |
| | 3795 $Atm^{-/-}$ | 800000 | 5 | 6.25 |
| | 3793 $Atm^{+/-}$ | 800000 | 0 | <1.25 |
| | 3835 $Atm^{-/-}$ | 800000 | 4 | 5.00 |
| | 3833 $Atm^{+/-}$ | 800000 | 0 | <1.25 |
| | 3834 $Atm^{+/+}$ | 800000 | 0 | <1.25 |
| | 3849 $Atm^{-/-}$ | 800000 | 3 | 3.75 |
| | 3850 $Atm^{+/-}$ | 800000 | 0 | <1.25 |
| | 3991 $Atm^{-/-}$ | 800000 | 2 | 2.50 |
| | 4286 $Atm^{-/-}$ | 800000 | 4 | 5.00 |
| | 4288 $Atm^{+/+}$ | 1600000 | 0 | <1.25 |
| | Total $Atm^{-/-}$ | 4800000 | 18 | 3.75 |
| | Total $Atm^{+/-}$; $Atm^{+/+}$ | 4800000 | 0 | <0.21 |

| | | | | |
|---|---|---|---|---|
| | 3777 *Atm*$^{-/-}$ | 800000 | 1 | 1.25 |
| | 3795 *Atm*$^{-/-}$ | 800000 | 3 | 3.75 |
| | 3793 *Atm*$^{+/-}$ | 800000 | 0 | <1.25 |
| | 3835 *Atm*$^{-/-}$ | 800000 | 4 | 5.00 |
| | 3833 *Atm*$^{+/-}$ | 800000 | 0 | <1.25 |
| Chr13 – 30 kb | 3834 *Atm*$^{+/+}$ | 800000 | 0 | <1.25 |
| | 3849 *Atm*$^{-/-}$ | 800000 | 2 | 2.50 |
| | 3850 *Atm*$^{+/-}$ | 800000 | 0 | <1.25 |
| | 3991 *Atm*$^{-/-}$ | 800000 | 3 | 3.75 |
| | 4286 *Atm*$^{-/-}$ | 800000 | 6 | 7.50 |
| | 4288 *Atm*$^{+/+}$ | 1600000 | 0 | <1.25 |
| | Total *Atm*$^{-/-}$ | 4800000 | 19 | 3.96 |
| | Total *Atm*$^{+/-}$; *Atm*$^{+/+}$ | 4800000 | 0 | <0.21 |

**Additional deletions (total 61 events in Atm$^{-/-}$) included in analyses in Fig. 2 and 6**

| | | | | |
|---|---|---|---|---|
| | 4286 *Atm*$^{-/-}$ | 800000 | 10 | 12.50 |
| Chr13 – 5 kb | 4286 *Atm*$^{-/-}$[b] | 800000 | 7 | 8.75 |
| | 4227 *Atm*$^{-/-}$[c] | 400000 | 3 | 7.50 |

Littermates are grouped.

[a] One deletion involving the $(TG)_n$ repeat excluded; see **Supplemental Fig. S1B**.

[b] From **Supplemental Table S6**.

[c] From **Supplemental Table S4**.

**Supplemental Table S3. Frequency of deletions in MRE11-proficient and -deficient testes**

| Hotspot pair | Mouse and genotype | No. haploid genomes | No. deletions | Frequency ($10^{-6}$) | No. PCR artifacts[a] |
|---|---|---|---|---|---|
| Chr13 – 5 kb | 63 *Mre11*$^{-/-}$ | 5600000 | 34 | 6.07 | 0 |
| | 62 *Mre11*$^{+/-}$ | 4800000 | 5 | 1.04 | 0 |
| | 127 *Mre11*$^{-/-}$ | 3200000 | 20 | 6.25 | 5 |
| | 128 *Mre11*$^{+/-}$ | 3200000 | 0 | <0.31 | 5 |
| | 194 *Mre11*$^{-/-}$[b] | 400000 | 3 | 7.50 | 0 |
| | Total *Mre11*$^{-/-}$ | 9200000 | 57 | 6.20 | 5 |
| | Total *Mre11*$^{+/-}$ | 8000000 | 5 | 0.63 | 5 |
| Chr13 – 15 kb | 63 *Mre11*$^{-/-}$ | 4000000 | 34 | 8.50 | 3 |
| | 62 *Mre11*$^{+/-}$ | 3200000 | 2 | 0.63 | 1 |
| | 4350 *Spo11*$^{-/-}$ | 4800000 | 0 | <0.21 | 1 |

Littermates are grouped.

[a] Excluded deletion events considered as PCR artifacts; see **Supplemental Fig. 1B**.

[b] From **Supplemental Table S4**.

**Supplemental Table S4. Frequency of deletions in *Mre11*$^{-/-}$ single and *Mre11*$^{-/-}$ *Tdp2*$^{-/-}$ double mutant testes.**

| Hotspot pair | Mouse and genotype | No. haploid genomes | No. deletions | Frequency ($10^{-6}$) | No. PCR artifacts[a] |
|---|---|---|---|---|---|
| Chr1 – 2 kb | 194 *Mre11*$^{-/-}$ | 2400000 | 13 | 5.42 | 0 |
| | 127 *Mre11*$^{-/-}$ | 2400000 | 27 | 11.25 | 0 |
| | 7 *Mre11*$^{-/-}$ *Tdp2*$^{-/-}$ | 2400000 | 9 | 3.75 | 0 |
| | 9 *Mre11*$^{-/-}$ *Tdp2*$^{-/-}$ | 2400000 | 11 | 4.58 | 0 |
| | Total *Mre11*$^{-/-}$ | 4800000 | 40 | 8.33 | 0 |
| | Total *Mre11*$^{-/-}$ *Tdp2*$^{-/-}$ | 4800000 | 20 | 4.17 | 0 |
| Chr13 – 5 kb | 4427 *Atm*$^{-/-}$ | 400000 | 3 | 7.50 | 0 |
| | 194 *Mre11*$^{-/-}$ | 400000 | 3 | 7.50 | 0 |
| | 7 *Mre11*$^{-/-}$ *Tdp2*$^{-/-}$ | 2800000 | 10 | 3.57 | 7 |
| | 9 *Mre11*$^{-/-}$ *Tdp2*$^{-/-}$ | 2800000 | 9 | 3.21 | 6 |
| | Total *Mre11*$^{-/-}$ *Tdp2*$^{-/-}$ | 5600000 | 19 | 3.39 | 13 |

[a] Excluded deletion events considered as PCR artifacts; see **Supplemental Fig. 1B**.

**Supplemental Table S5. Frequency of deletions in *Atm*$^{-/-}$ single and *Atm*$^{-/-}$ *Tdp2*$^{-/-}$ double mutant testes**

| Hotspot pair | Mouse and genotype | No. haploid genomes | No. deletions | Frequency ($10^{-6}$) | No. PCR artifacts[a] |
|---|---|---|---|---|---|
| Chr1 – 2 kb | 3962 *Atm*$^{-/-}$ *Tdp2*$^{+/-}$ | 3200000 | 20 | 6.25 | 0 |
| | 3963 *Atm*$^{-/-}$ *Tdp2*$^{-/-}$ | 3200000 | 21 | 6.56 | 0 |
| | 4369 *Atm*$^{-/-}$ *Tdp2*$^{+/-}$ | 2400000 | 23 | 9.58 | 0 |
| | 4370 *Atm*$^{-/-}$ *Tdp2*$^{-/-}$ | 5600000 | 9 | 1.61 | 0 |
| | Total *Atm*$^{-/-}$ *Tdp2*$^{+/-}$ | 5600000 | 43 | 7.68 | 0 |
| | Total *Atm*$^{-/-}$ *Tdp2*$^{-/-}$ | 8800000 | 30 | 3.41 | 0 |
| Chr13 – 5 kb | 3962 *Atm*$^{-/-}$ *Tdp2*$^{+/-}$ | 1600000 | 3 | 1.88 | 0 |
| | 3963 *Atm*$^{-/-}$ *Tdp2*$^{-/-}$ | 1600000 | 2 | 1.25 | 0 |
| | 4369 *Atm*$^{-/-}$ *Tdp2*$^{+/-}$ | 4800000 | 29 | 6.04 | 2 |
| | 4370 *Atm*$^{-/-}$ *Tdp2*$^{-/-}$ | 4800000 | 7 | 1.46 | 1 |
| | 4225 *Atm*$^{-/-}$ *Tdp2*$^{-/-}$ | 3200000 | 13 | 4.06 | 1 |
| | Total *Atm*$^{-/-}$ *Tdp2*$^{+/-}$ | 6400000 | 32 | 5.00 | 2 |
| | Total *Atm*$^{-/-}$ *Tdp2*$^{-/-}$ | 9600000 | 22 | 2.30 | 2 |

Littermates are grouped.
[a] Excluded deletion events considered as PCR artifacts; see **Supplemental Fig. 1B**.

**Supplemental Table S6. Frequency of deletions in MRE11-proficient and *Mre11$^{H129N}$* testes**

| Hotspot pair | Mouse and genotype | No. haploid genomes | No. deletions | Frequency ($10^{-6}$) |
|---|---|---|---|---|
| Chr13 – 5 kb | 102 *Mre11$^{H129N/–}$* | 2800000 | 2 | 0.71 |
| | 103 *Mre11$^{H129N/–}$* | 2800000 | 2 | 0.71 |
| | 104 *Mre11$^{+/–}$* | 4800000 | 3 | 0.63 |
| | 4286 *Atm$^{–/–}$* | 800000 | 7 | 8.75 |
| | Total *Mre11$^{H129N/–}$* | 5600000 | 4 | 0.71 |

Littermates are grouped.

**Supplemental Table S7. Frequency of deletions in NBS1-proficient and *Nbs1*$^{\Delta B}$ testes and sperm**

| Hotspot pair | Tissue | Mouse and genotype | No. haploid genomes | No. deletions | Frequency ($10^{-6}$) |
|---|---|---|---|---|---|
| Chr13 – 5 kb | sperm | 95 *Nbs1*$^{\Delta B/\Delta B}$ | 1600000 | 4 | 2.50 |
| | | 96 *Nbs1*$^{\Delta B/\Delta B}$ | 1600000 | 1 | 0.63 |
| | | 4229 *Nbs1*$^{\Delta B/\Delta B}$ | 1600000 | 1 | 0.63 |
| | | 4230 *Nbs1*$^{\Delta B/+}$ | 1600000 | 0 | <0.63 |
| | | 4292 *Nbs1*$^{\Delta B/\Delta B}$ | 2400000 | 3 | 1.25 |
| | | 4293 *Nbs1*$^{\Delta B/\Delta B}$ | 800000 | 0 | <0.13 |
| | | 4290 *Nbs1*$^{+/+}$ | 4000000 | 3 | 0.75 |
| | | 4291 *Nbs1*$^{+/+}$ | 4000000 | 4 | 1.00 |
| | | Total *Nbs1*$^{\Delta B/\Delta B}$ | 8000000 | 9 | 1.13 |
| | | Total *Nbs1*$^{\Delta B/+}$; *Nbs1*$^{+/+}$ | 9600000 | 7 | 0.73 |
| | testis | 95 *Nbs1*$^{\Delta B/\Delta B}$ | 2400000 | 2 | 0.83 |
| | | 96 *Nbs1*$^{\Delta B/\Delta B}$ | 2400000 | 0 | <0.42 |
| | | 4154 *Nbs1*$^{\Delta B/\Delta B}$ | 1600000 | 0 | <0.63 |
| | | 4153 *Nbs1*$^{\Delta B/+}$ | 1600000 | 1 | 0.63 |
| | | Total *Nbs1*$^{\Delta B/\Delta B}$ | 6400000 | 2 | 0.31 |
| Chr19 – 2 kb | sperm | 4279 *Atm*$^{+/-}$ | 800000 | 0 | <1.25 |
| | | 4280 *Atm*$^{+/+}$ | 800000 | 1 | 1.25 |
| | | 4229 *Nbs1*$^{\Delta B/\Delta B}$ | 2400000 | 0 | <0.42 |
| | | 4230 *Nbs1*$^{\Delta B/+}$ | 2400000 | 2 | 0.83 |
| | | 4292 *Nbs1*$^{\Delta B/\Delta B}$ | 4000000 | 5 | 1.25 |
| | | 4290 *Nbs1*$^{+/+}$ | 4000000 | 2 | 0.50 |
| | | Total *Nbs1*$^{\Delta B/\Delta B}$ | 6400000 | 5 | 0.78 |
| | | Total *Nbs1*$^{\Delta B/+}$; *Nbs1*$^{+/+}$ | 6400000 | 4 | 0.63 |
| | testis | 4154 *Nbs1*$^{\Delta B/\Delta B}$ | 800000 | 1 | 1.25 |
| | | 4153 *Nbs1*$^{\Delta B/+}$ | 800000 | 2 | 2.50 |

Littermates are grouped.

**Supplemental Table S8. Frequency of microdeletions in individual *Mre11*$^{-/-}$ and *Atm*$^{-/-}$ amplicons**

| Genotype | Amplicon | No. reads | No. reads with deletions | Unique deletions[a] | Frequency of reads with deletions ($10^{-6}$) |
|---|---|---|---|---|---|
| *Mre11*$^{-/-}$ | 1 | 3448619 | 486 | 176 | 140.93 |
| | 2 | 6019263 | 1063 | 242 | 176.60 |
| | 3 | 5316448 | 956 | 204 | 179.82 |
| | 4 | 6133303 | 812 | 182 | 132.39 |
| | Total | 20917633 | 3317 | 804 | 158.57 |
| *Atm*$^{-/-}$ | 1 | 3242455 | 162 | 68 | 49.96 |
| | 2 | 2462674 | 145 | 77 | 58.88 |
| | 3 | 3261329 | 224 | 105 | 68.68 |
| | 4 | 2817262 | 116 | 59 | 41.17 |
| | Total | 11783720 | 647 | 309 | 54.91 |

[a] Total unique deletions obtained in that amplicon; note that the same deletion can be obtained independently in the overlapping amplicon.
